# Pre-training Genomic Language Model with Variants for Better Modeling Functional Genomics

**DOI:** 10.1101/2025.02.26.640468

**Authors:** Tianyu Liu, Xiangyu Zhang, Jiecong Lin, Luca Pinello, Rex Ying, Hongyu Zhao

## Abstract

Sequence-to-function models can predict gene expression from sequence data and be used to link genetic information with transcriptomics data to understand regulatory processes and their effects on complex phenotypes. The genomic language models are pre-trained with large-scale DNA sequences and can generate robust representations of these sequences by learning the genomic context. However, few studies can estimate the predictability of gene expression levels and bridge these two classes of models together to explore individualized gene expression prediction. In this manuscript, we propose UKBioBERT as a DNA language model pre-trained with genetic variants from UK BioBank. We demonstrate that UKBioBERT generates informative embeddings capable of identifying gene functions, and improving gene expression prediction in cell lines, thereby enhancing our understanding of gene expression predictability. Building upon these embeddings, we combine UKBioBERT with state-of-the-art sequence-to-function architectures, Enformer and Borzoi, to create UKBioFormer and UKBioZoi. These models exhibit better performance in predicting highly predictable gene expression levels and can be generalized across different cohorts. Furthermore, UKBioFormer effectively captures the relationship between genetic variants and expression variations, enabling in-silico mutation analyses and eQTL identification. Collectively, our findings underscore the value of integrating genomic language models and sequence-to-function approaches for advancing functional genomics research.

## 1 Introduction

Deciphering DNA as a language is a challenging but essential task to understand the complexity in genetics and genomics [1–8]. Previous research has demonstrated that the functional elements in genomes, such as cis-regulatory elements (CREs) and trans-regulatory elements, may be affected by many factors including distance to targeted genes as well as the context of their nearby genomics regions [9, 10]. These properties suggest the existence of polysemy and distant semantic relationships in DNA sequence, which are also found in natural language [11]. Therefore, modeling DNA as a language of life can also be based on the advanced techniques in Natural Language Processing (NLP) [12], and many foundation models based on the architecture of Transformer [13] have been proposed to model DNA sequences, and led to better understanding of functional genomics in different species [14] or cell lines [15].

There are two major types of genomic language models (gLMs), either learning the context of DNA sequences by masked language modeling (MLM) or learning the causal relationship in nucleotide sequences by causal language modeling (CLM). The representative base model for the first class is the bidirectional transformer (BERT) [16]. For example, DNABERT [11], DNABERT-2 [17], and DNABERT-S [14] utilize BERT as the base model and pre-train it with DNA sequences from the reference genome [18]. Furthermore, GENA-LM [19] pre-trains the base BERT model with both reference genome and single nucleotide polymorphisms (SNPs) from the 1000 Genomes Project [20], and provides models with different scales for users. Training the model based on BERT can enhance our understanding of the context information of one DNA sequence and thus it will generate better presentations for applications, in principle. The representative base model for the second class is a generative pre-trained transformer (GPT) [21]. As an example, DNAGPT [22] utilizes GPT-based models for pre-training and it can generate new DNA sequences. HyenaDNA [2] and Evo [23] are based on the Hyena convolution network [24], but their pre-training strategies are CLM, so they are also focusing on generating new DNA sequences or other large molecules (e.g., RNA or amino acid sequences).

Despite the exciting progress made by gLMs, there are concerns about applying these foundation models in genomic research. First, CLM-based DNA language models suffer from hallucination when generating new DNA sequences [25], and are not good at modeling the genomic information with the consideration of context, as shown in the recent benchmarking analysis of DNA sequence embeddings from various models [26].

Second, although MLM-based DNA language models are better candidates for learning the context information as well as encoding sequence information into the embedding space, the reliability of generated embeddings is affected by multiple factors, including the quality of pre-training datasets, the choice of pre-training task, the evaluation metrics, and the context window size, can all affect the model performance [26, 27]. Moreover, gLMs such as GPN [28] and GENA-LM consider including SNPs [29] to enhance model performance based on the principle of data augmentation, but there is no demonstration of the necessity of using variants versus using reference genome with the random augmentation approach [30]. Furthermore, the downstream applications considered by these methods are still based on reference genome sequences, and thus the pre-training data have a weak link with real application scenarios. These settings may limit the contribution of introducing human variants in model pre-training, although it is well-known that many genetic variants play an important role in genetic regulation, disease onset, and progression [31, 32].

To better model DNA as a language as well as address the limitations of the existing methods, we propose UKBioBERT, a foundation model pre-trained with both reference genome and variant information from UK BioBank [33]. We gather variants from approximately 300,000 UK Biobank participants with European ancestry and select the best approach to leveraging the advantages of pre-trained weights from other gLMs to formalize our models efficiently. We propose a new metric to evaluate the quality of gLMs by paying attention to the ability of a model to distinguish sequences with different functions. Furthermore, we investigate the contribution of genetic variants by considering downstream applications including cell-type-specific gene expression prediction and individualized gene expression prediction, which are more related to sequence-to-function analysis. The representations of UKBioBERT improved over baseline models, and explain the difficulties of expression prediction across individuals and cohorts. We also demonstrate that gene expression prediction at the individual level can be further improved by fine-tuning Enformer [30] with individual genome information and embeddings from UKBioBERT, denoted as UKBioFormer. Finally, we analyze the interpretability of the embeddings from UKBioBERT and identify both common and rare variant effects validated by expression quantitative trait loci (eQTL) information based on UKBioFormer. Overall, UKBioBERT and UKBioFormer offer useful tools for researchers to explore the power of variant modeling with sequences from different population-level datasets.

## 2 Results

### Method overview

The development of UKBioBERT is inspired by the training process of GPN [28] and Gena-LM [19], as well as data augmentation from deep learning [34]. We pre-trained UKBioBERT based on the checkpoint from DNABERT2 [17] and genomic information with variants from UK BioBank with a self-supervised learning framework. Recent benchmarking analyses demonstrated that DNABERT2 performed well for several gene-centric tasks, such as gene finding, variant effect prediction, and gene expression prediction [35–37]. These observations suggest the usefulness of modeling genomic language with the awareness of frequency and context differences contributed by variants. The embeddings from UKBioBERT can help state-of-the-art (SOTA) method EPInformer [15] for better-predicting gene expression captured by different techniques at the cell-type level. These contributions are summarized in Figure 1 (a). Moreover, the embeddings generated from individual genomic information can also explain the predictability of gene expression levels from a novel angle. The predictability of one gene is defined as the accuracy of predicted gene expression based on DNA sequences only. Furthermore, we selected the optimal models from a collection of UKBioBERT and combined the embeddings with Enformer to fine-tune the fused model UKBioFormer for gene expression prediction and eQTL identification at the individual level. These contributions are summarized in Figure 1 (b). Details of our selected tools are further explained in the Methods section.

**Fig. 1.**
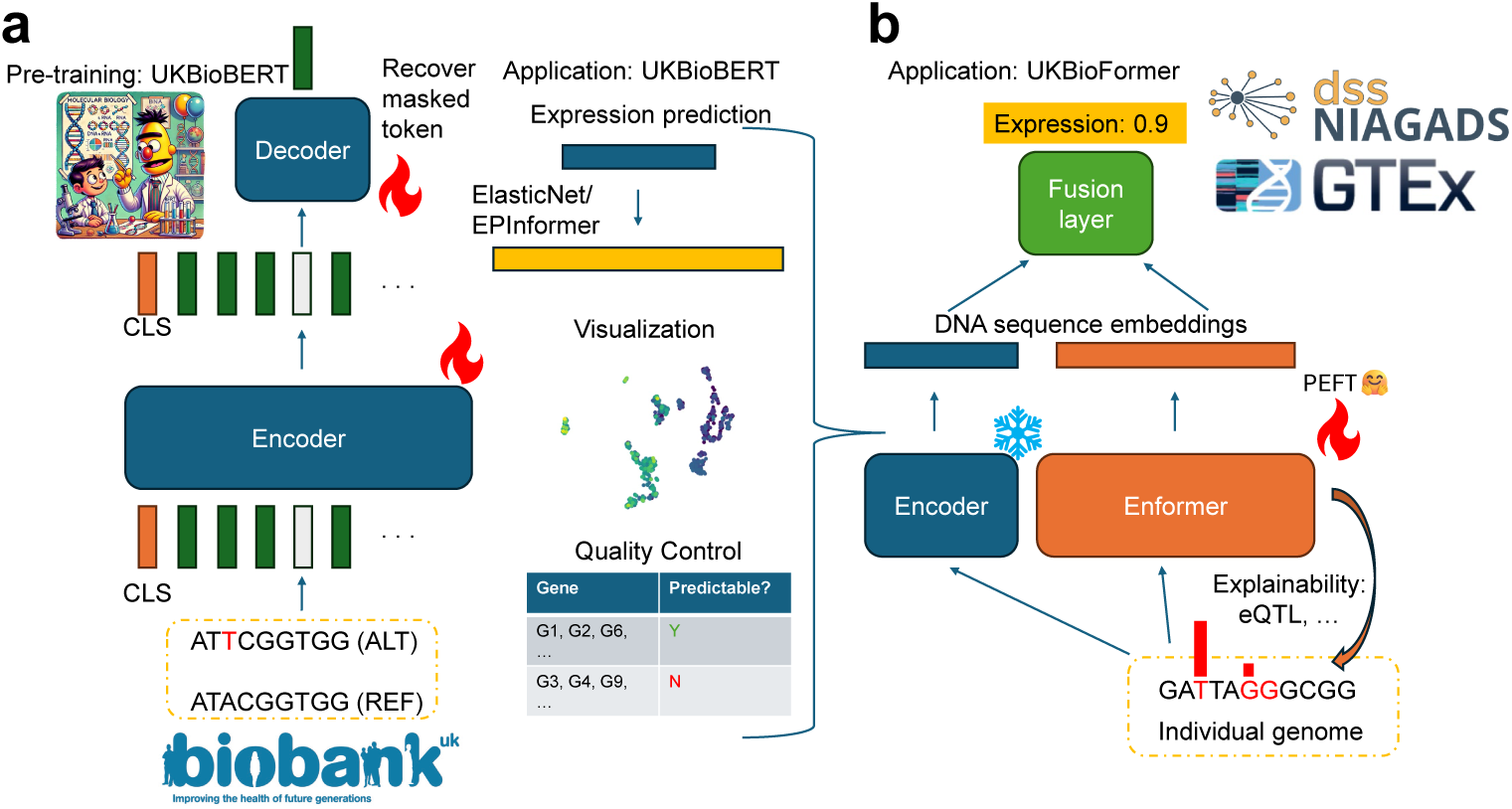
The landscape of UKBioBERT and UKBioFormer. The fire logo represents the adjustment of model parameters, and the ice logo represents the freezing of model parameters. The logos of UK BioBank, GTEx, and ROSMAP come from the official websites. (a) The pre-training stage and related downstream applications of UKBioBERT. We utilize masked language modeling to pre-train UKBioBERT using modified reference (REF) genome (ALT). Our sequence starts with the Classify token (CLS). The cartoon logo is generated by GPT-4o [38]. (b) The fine-tuning stage and related downstream applications of UKBioFormer. We fuse Enformer and UKBioBERT to enhance gene expression prediction, and also leverage Parameter Efficient Fine-Tuning (PEFT) for efficient training.

### Pre-trained UKBioBERT learns better gene representations

Previous genomic language models evaluated their pre-training stage based on the validation loss and further investigated the expressive power of their pre-trained model based on downstream applications of the DNA sequence embeddings, such as enhancer identification and species classification [17, 35]. However, these tasks did not consider an important problem in the pre-training stage of foundation models, known as model collapse or context collapse. Model collapse is defined as that the pre-trained model cannot distinguish between data that differ in input features. Furthermore, this problem is also reflected in the embedding space of low-performance models, that is, the embeddings are similar for DNA sequences containing different contexts. Since the identity of a given DNA sequence can be varied, and we mainly consider gene expression levels, we focus on the function of genes as the main labels. The function of genes is known to be highly associated with expression levels, and thus gene function is a suitable label to describe the level of model collapse. Specifically, we compared gLMs by using different models to embed DNA sequences with different functional labels and tested the clustering performance based on these labels. To make a fair comparison, we subset the DNA sequences of human genes from a reference genome and pair the sequences with the corresponding functional labels from [39], as inputs and observed labels. We considered Normalized Mutual Information (NMI), Adjusted Rand Index (ARI), and Average Silhouette Width (ASW) as evaluation metrics [40, 41] and averaged them as a score to compare different gLMs, with a higher average score representing a better model. We selected gLMs (with one LLM) [2, 11, 17, 19, 30, 42–51] from review and benchmarking papers detailed in the Methods section with UKBioBERT based on both scores and model scales. We also compared the embeddings generated from UKBioBERT with those from pre-trained DNABERT2 as the base model, those with random initialization, those with contrastive learning pre-training [52, 53], and those with LD score regression pre-training, shown in Extended Data Figure 1 (a). According to this figure, choosing a base model trained with a reference genome is helpful in learning a good gene representation, and MLM is best for pre-training. One possible reason is that the masked modeling approach is closer to the biological interpretation of variants, rather than statistical-based or computational-based methods such as LD score modeling and contrastive learning. We follow the default pooling approaches from these methods to generate the embeddings, while the embeddings from UKBioBERT benchmarked in this section are generated based on the embeddings of the CLS token. Details of the comparison process are explained in the Methods section.

Our results are shown in Figures 2 (a)-(d). The achievement of UKBioBERT can be attributed to the richness of the training dataset, whose statistics are shown in Figure 2 (a) and we utilized over 13 million variants for the continued pre-training stage. We also presented the Avg score vs. training step curve in Extended Data Figure 2 (a), which shows that the optimal checkpoint is obtained at the early stage of pre-training, and the reduction of clustering score can be treated as a signal of overfitting. Finally, we explored the data scaling laws in the pre-training stage to investigate if including more variants can improve the quality of gene representations generated by UKBioBERT, shown in Figure 2 (b). In the same figure, we found that increasing the proportion of used variants could also introduce more variants near TSS or gene body for model training, which might explain the improvement of gene functional annotation performance. According to this figure, incorporating more human variants can improve the identification of gene functions and thus the quality of representations will improve as data scale increases based on UK BioBank data. However, we also intend to maintain a balance between the pre-training quality and the ability to predict gene expression. Therefore, we further evaluated the performances of using gene embeddings to predict individual gene expression levels as well as the quality of the embedding based on DNABERT2 pre-trained with gnomAD [54] (a dataset contains 0.7 billion human variants), and the results are summarized in Extended Data Figure 1 (a) and Extended Data Figure 2 (b). According to Figure 2 (c), UKBioBERT has the highest average score in the representation of the genes with their corresponding functions, without including the functional labels in the model pre-training stage. Furthermore, UKBioBERT has a relatively feasible parameter size compared with billion-level models, which improved its efficiency. UKBioBERT can also encode DNA sequences with arbitrary lengths by automatically dealing with the length of the input sequence. Moreover, we found that fine-tuning LLMs with DNA sequences and taking the embeddings do not help the model learn the functional information of genes, demonstrated by the low score of embeddings from Llama 3.1 7B [49, 50]. Models pre-trained with gene expression information did not perform well in this case, and one explanation could be that their attention mechanism assigned higher weights for the transcription starting site (TSS) rather than other regions, while the other explanation is that the random data augmentation used in the pre-training process is not as good as variant-based data augmentation. To visualize the differences of gene embeddings, we further show the sequence embeddings from UKBioBERT and other top baselines using uniform manifold approximation and projection (UMAP) [55] in Figure 2 (d). According to this figure, the panel corresponding to UKBioBERT clearly identifies the protein-encoding genes (with the pink labels) and other genes with different functions with specific clusters. Based on these figures, our current setting has already optimized the representations of genes for both the gene function separation task and the expression prediction task. Therefore, we demonstrated the contribution of incorporating human variants into the model pre-training stage within a specific range of data contexts. The UMAPs for all baselines as well as the meaning of annotations are summarized in Extended Data Figures 1 (b) and (c). Therefore, the embeddings from UKBioBERT have the best performance in uncovering the gene functions directly from the sequence information, and thus can be treated as good representations with the preservation of genomic context.

**Fig. 2.**
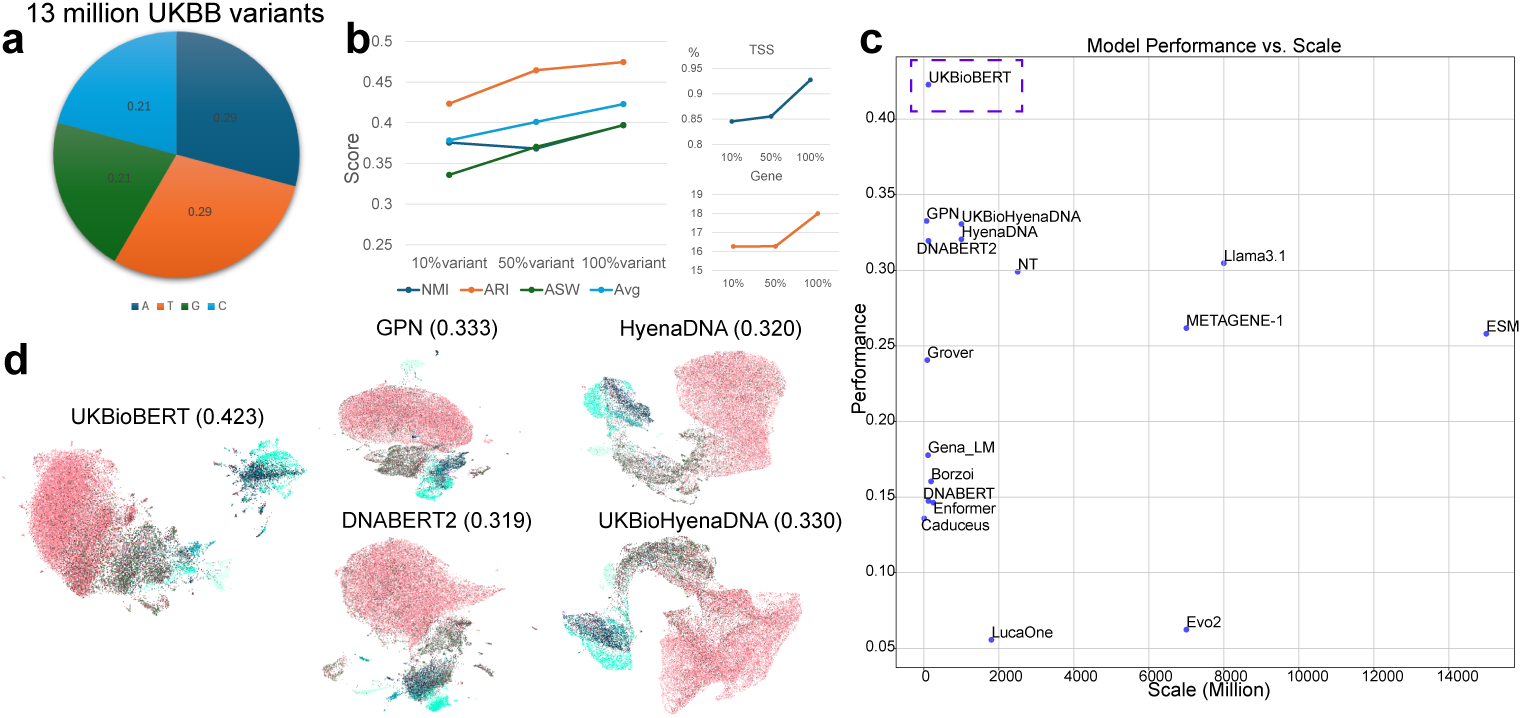
Main results of model pre-training. (a) The statistics of the human variant corpus used for pre-training. (b) The relationship between model performance and factors. The left panel shows the relationship between proportion of used variants and clustering performance, while the right panel shows the relationship between the proportion of variants near TSS and the proportion of used variants, and the relationship between the proportion of variants in gene body and the proportion of used variants. The gene functional annotation list can be found in Extended Data Figure 1 (b). (c) The scatter plot for the relationship between clustering performances and model scales. We highlight the best model UKBioBERT. (d) The UMAP visualization of UKBioBERT and other baselines with top performances, colored by gene types. The average clustering scores are annotated with UMAP panels.

### Embeddings from UKBioBERT can enhance model performance for gene expression prediction at the cell-type level

Transformer-based methods have been widely shown to effectively predict gene expression levels from sequences. Furthermore, recent research argued that incorporating and modeling external information, such as the interaction between promoter and potential enhancers, epigenetic signals, as well as chromatin contact data, can further improve the model’s performance in predicting gene expression measured by different techniques, for example, CAGE-seq and RNA-seq [15]. The authors of EPInformer demonstrated the advantages of EPInformer versus other transformer-based genomic deep-learning models in predicting gene expression levels by training genes on different chromosomes. Therefore, we extended this concept to a broadened domain and demonstrated that we could incorporate more informative signals to enhance EPInformer, including the DNA sequence embeddings from UKBioBERT and EpiBERT [56], as well as the gene description embeddings from scELMo [57]. The concept of DNA sequence embedding was discussed in our previous section, where our sequence embeddings allowed us to capture the functional information in the corresponding genomic region under the awareness of the whole DNA sequence library. Moreover, we could also utilize Large Language Models to generate and encode the descriptions of gene functions, and such text-based gene embeddings can also introduce the numerical representation of gene functions to help predict the gene expression profiles. To incorporate gene embeddings as new inputs, we concatenated the original sequence embeddings (promoters and enhancers), epigenetic embeddings, and newly-added gene embeddings, and then passed the joint embeddings to the prediction head, to make the final prediction. Therefore, we improved EPInformer by modeling the default genomic information, the sequence embeddings from promoter and enhancers, as well as text-based gene embeddings, to formulate a new model and train it based on the same datasets. Due to data access limitations, we focused only on human cell lines with available promoter and enhancer information and validated our model with the corresponding CAGE-seq and RNA-seq results. We considered K562, GM12878, and HepG2 cell lines, where we can retrieve enough information to implement the default mode of EPInformer. We also considered modeling silencers under the same framework, but there was no improvement after including the silencer information [58], shown in Extended Data Figure 3. Our hyper-parameter setting and data splits (12-fold cross-validation) follow the same settings from EPInformer, and thus our results can directly reveal the performance difference.

In Figure 3 (a), we visualize the comparisons among different modes of EPInformer, including the mode without pre-trained convolution network for identifying the enhancers (A), EPInformer with DNA sequence embeddings from EpiBERT (B), the default mode (C), EPInformer with DNA sequence embeddings generated by Hye-naDNA [2] (D), EPInformer with DNA sequence embeddings from UKBioBERT (E), EPInformer with text-based embeddings from scELMo (F), EPInformer with embeddings from UKBioBERT and embeddings from scELMo (G). According to this figure (each dot represents the Pearson Correlation Coefficient (PCC) between predicted and observed values evaluated), the joint mode G outperformed the default mode of EPInformer, and also improved the method compared with other ablation versions. Compared with other embeddings, using EpiBERT could not improve the performance of EPInformer. For the testing results based on CAGE-seq from GM12878, our method had an average PCC over 0.9, significantly outperforming other methods. Furthermore, our method also had a lower variance in 3 out of 4 datasets compared with the raw mode, which suggests that the joint mode has better stability. To further strengthen our conclusion, we utilized RNA-seq data from HepG2 and applied EPInformer for predicting gene expression levels. According to Extended Data Figure 4 (a), using embeddings from scELMo and UKBioBERT outperforms other choices again, and thus the generalization ability of our proposed method is guaranteed.

**Fig. 3.**
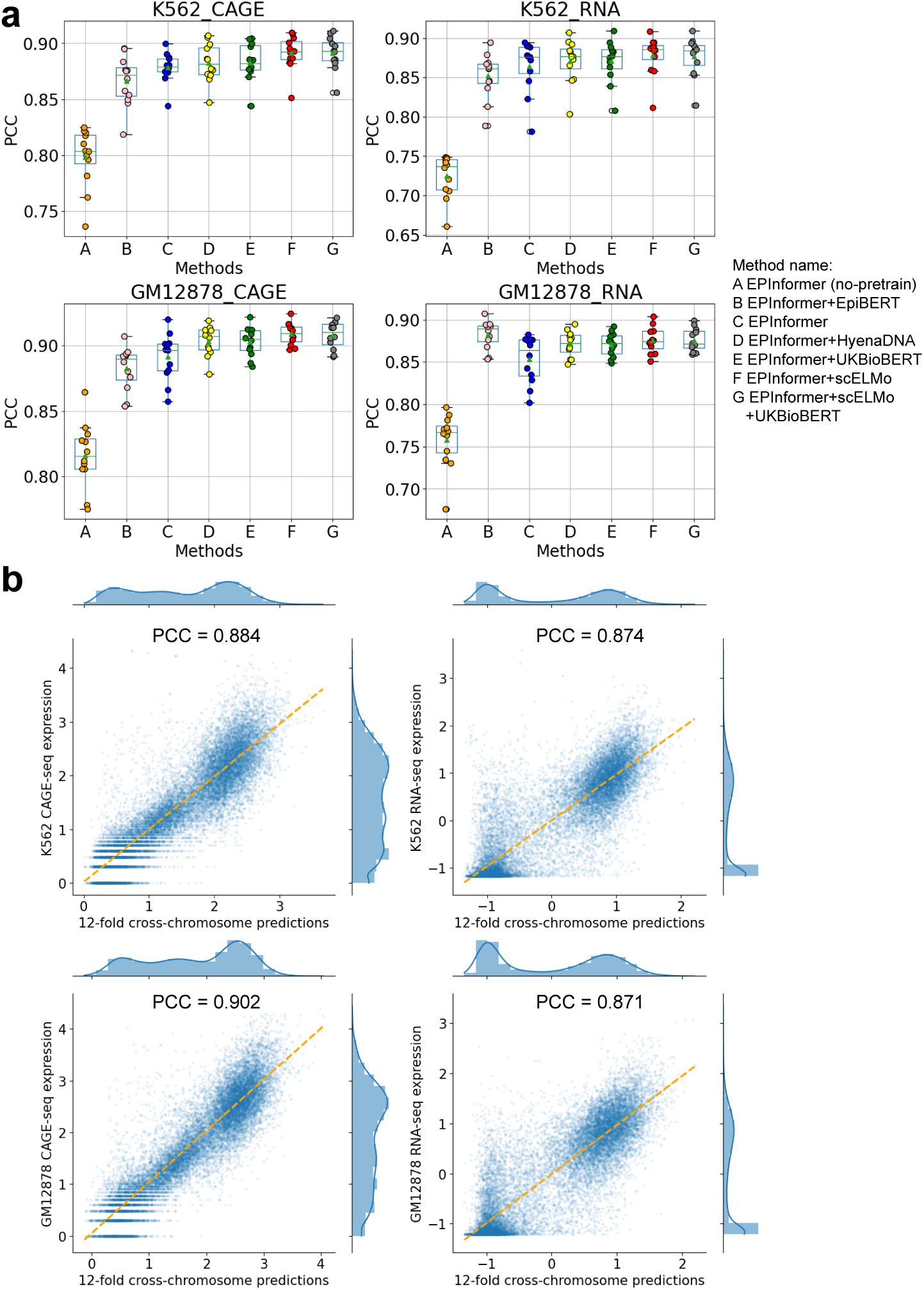
The results of optimizing gene expression prediction in the K562 and GM12878 cell lines with UKBioBERT. (a) Comparisons of different methods for gene expression prediction across techniques and cell lines. The metric used in this section is PCC. (b) The scatter plot and distribution plot of the best performer (combination E) for the relationship between observed expression and predicted expression. The joint PCC is annotated and the yellow line is fit with two sets of expression levels.

To further investigate the properties of model outputs, we visualize the predicted and observed gene expression levels from these 12 folds in Figure 3 (b) and Extended Data Figure 4 (b), as well as their distributions. These figures are annotated with a set of overall PCCs, which are all higher than 0.8. Moreover, the predicted gene expression levels show a smoother distribution, especially in the prediction of RNA-seq data, which also suggests that the RNA-seq data might contain noise which could be reduced by our method.

### Embeddings from UKBioBERT can recover the predictability of gene expression levels across different conditions

Since UKBioBERT is pre-trained with variants from large-scale population-level data, we also assumed that the DNA sequence embeddings from UKBioBERT can help us predict the gene expression profiles at the individual level, and further help explain the predictability of specific genes. Here we considered baselines including the traditional method ElasticNet [59], the pre-trained model with zero-shot learning mode Enformer, and fine-tuned Enformer based on personalized data, known as Personalized Enformer (Performer) [7]. To predict gene expression levels, we used the embeddings from UKBioBERT as inputs and ElasticNet as the predictor for evaluation, and chose 41 representative genes for analysis. These genes were either considered in previous research [7] or have gene expression level heritability estimated based on Linear Mixed Models (LMMs) [60]. In this study, we collected 670 samples from GTEx data with paired RNA-seq and WGS data, and performed 5-fold cross-validation to report the testing performances measured by three metrics: Pearson Correlation Coefficient (PCC), *R*^2^ score (R2), and mean squared error (MSE).

The PCC between the predicted and observed expression levels is shown in Figure 4 (a). According to this figure, ElasticNet, UKBioBERT, and Performer all outperformed Enformer, suggesting that the zero-shot learning mode of Enformer cannot directly predict individualized gene expression levels well. Furthermore, these three methods showed similar results, supported by the strong correlation of different scores (PCC=0.988, p-value=2.19e-34 between UKBioBERT and ElasticNet, and PCC=0.991, p-value=1.92e-36 between UKBioBERT and Performer). We found that this is not only reflected in the results of PCC for the two groups of genes but also shown by studies on the R2 score and MSE, shown in Extended Data Figures 5 (a)-(d). On average, ElasticNet had slightly better performance, similar to the conclusion from a previous study [7]. However, we also found that no model can perfectly predict the expression of all genes and there is a significant difference in the prediction performance across different genes. We first tried to use heritability to explain this observation, but Figure 4 (b) showed that the heritability of the selected gene group did not have an apparent correlation with prediction performance. We further computed the correlation between metrics and heritability based on both PCC and Spearman Correlation Coefficient (SCC), but both results did not show significant correlation (p-value=0.19 for PCC, p-value=0.17 for SCC), shown in the left two panels of Figure 4 (c). Therefore, gene heritability does not explain the prediction differences across genes. Interestingly, by visualizing the relationship between the gene embeddings from UKBioBERT and expression levels, we found that the individuals with similar expression patterns are more likely to be clustered in one group, especially for genes with good prediction performance, shown in Extended Data Figure 7. Therefore, the clustering performance of individuals based on expression levels may be a good explanation for gene predictability. We calculated the clustering metrics based on NMI for all 41 genes and visualized the strong relationship between prediction performances and clustering metrics in the middle panels of Figure 4 (c), which supports our inference. To investigate biological reasons for this strong relationship, we also analyzed the correlation between PCCs and other genome-level factors, such as gene location, number of unique DNA sequences, number of enhancers/promoters, and gene function score (GIFts) from GeneCard [61, 62], as discussed in Appendix A. Based on our analysis, the GIFts scores have a significant negative correlation with prediction performance (Extended Data Figure 12 (e)). It is also difficult to predict expression levels for genes with more functions because multiple gene functions cannot be fully explained only by sequence information. Therefore, before starting prediction, we recommend visualizing such relationships for the training datasets in low-dimensional space such as UMAP, computing the clustering scores, and searching the GIFts score to determine whether the specific gene may be predicted from sequences or not.

**Fig. 4.**
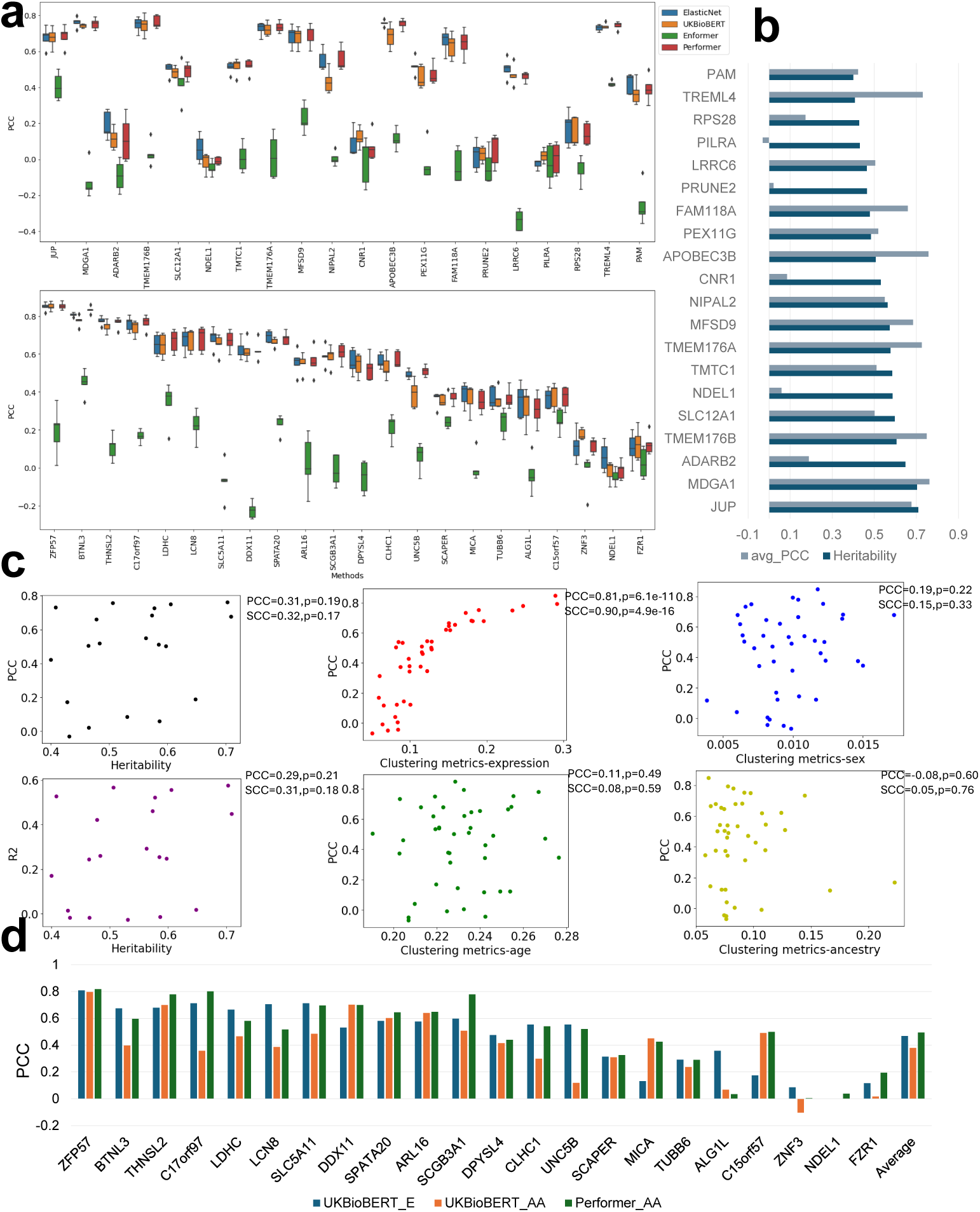
Performance comparisons of gene expression prediction among UKBioBERT and selected baselines. (a) Comparison of PCC across two sets of genes for the selected methods. (b) Relationship between averaged PCC across five folds and heritability for selected genes. (c) Panel list for the relationship between scores of metrics and metadata information of individuals in GTEx. The metrics include PCC and R2, while the metadata includes heritability, cluster-based expression levels, age, sex, and ancestry. Each dot represents one gene. (d) Comparison of PCC for models tested with individuals from different ancestries across selected genes. The notation “E” represents results tested based on people with European ancestries. The notation “AA” represents results tested based on people with African American ancestries.

The gene embeddings from UKBioBERT can also be used to analyze the relationship between prediction performance and other metadata or environmental factors, such as age, gender, and ancestry. We also visualize such relationships between the PCC and these three factors in the middle and right panels of Figure 4 (c), showing that the accuracy of gene expression prediction at the population level is not strongly correlated with the above factors. The prediction performance is only related to how easily the corresponding gene sequences can be distinguished, and it is biased to attribute the expression of a specific gene to factors such as ancestry.

Moreover, we tested different pooling methods to generate the gene embeddings based on UKBioBERT for gene expression prediction, including CLS token representation (CLS), mean pooling representation (Mean), and max pooling representation (Max). We performed tests based on the top five genes ranked by PCCs and our results implied that the Mean pooling and Max pooling are comparable, and they both surpassed the CLS pooling approach, shown in Extended Data Figure 8 (a). This observation is consistent with previous analysis of using LLM embeddings for regression problems in medical applications [63]. We also explored possible data-efficient learning designs for predicting gene expression levels with the gene ZFP57, which has the best prediction performance, and model UKBioBERT. We subsampled the training datasets with different proportions of individuals and tested the prediction performances, shown in Extended Data Figure 8 (b). According to this figure, using all the samples had the best performance (highest average PCC). Furthermore, Extended Data Figure 8 (c) demonstrates that using the full genome information from both two parents can generate better prediction performance than using partial information. Therefore, our current implementation is optimal at the data level and we suggest including as many samples as possible for better prediction.

### Fine-tuned UKBioFormer predicts more accurate gene expression levels across individuals compared with baselines

The previous section mainly focused on gene expression predictability by the representations from UKBioBERT, which did not cover two major challenges of personalized gene expression prediction: cross-cohort expression prediction and group-level gene expression prediction. The cross-cohort expression prediction focuses on predicting gene expression for unseen cohort-level data, such as sequence data from individuals in a new study or of different ancestry [59]. The group-level gene expression focuses on extending the training design of prediction at the single-gene level to multiple genes which belong to the same pathway or share the same enhancers, starting from genes pre-trained with Enformer. If the model can learn the synergetic effect and shared regulation across different genes, the performance of expression prediction might be improved. Training a model for multi-gene prediction also reduces cost. In addition to the four baseline models presented previously, we considered several new models for cross-individual prediction, denoted as UKBioFormer, UKBioZoi, Gena LM, HyenaDNA, and Basenji2 [64]. UKBioFormer integrates Enformer and UKBioBERT for more accurate predictions with a parameter-efficient-fine-tuning (PEFT) design, and UKBioZoi follows the same idea but the base model is Borzoi. HyenaDNA and genaLM are two gLMs trained with ElasticNet for expression prediction, and Basenji2 is a supervised model designed to predict expression profiles across different conditions from sequences. As shown in Figure 1 (a), we concatenated the sequence embeddings generated from Enformer or Borzoi with gene embeddings from UKBioBERT as a new gene representation, and passed the representation to the prediction head and produced the individual-specific gene expression. We used the same metrics to evaluate model performances.

To perform a systematical evaluation, we utilized the same two sets of genes and visualized the prediction metrics in Figures 5 (a), (b), Extended Data Figures 5 (a)-(d), and Extended Data Figures 6 (a) and (b). According to these figures, UKBioFormer had better performance than Performer and UKBioZoi on average, and its performance is consistent with our previous analysis based on gene predictability. If we selected genes with good predictability (PCC larger than 0.6), UKBioFormer had better performances than Performer in 63.3% genes. Furthermore, UKBioFormer had more genes with improvement over Performer, shown in Extended Data Figure 6 with higher x-axis value. Furthermore, UKBioFormer had a more efficient design as its running time and GPU memory consumption are lower than Performer and ElasticNet, shown in Extended Data Figures 9 (a)-(c). Although UKBioZoi had a slightly worse performance, it has shorter training time and less memory cost. Therefore, we recommend using UKBioFormer first for personalized gene expression prediction, and if the computation resource becomes a limitation, using UKBioZoi as an alternative plan.

**Fig. 5.**
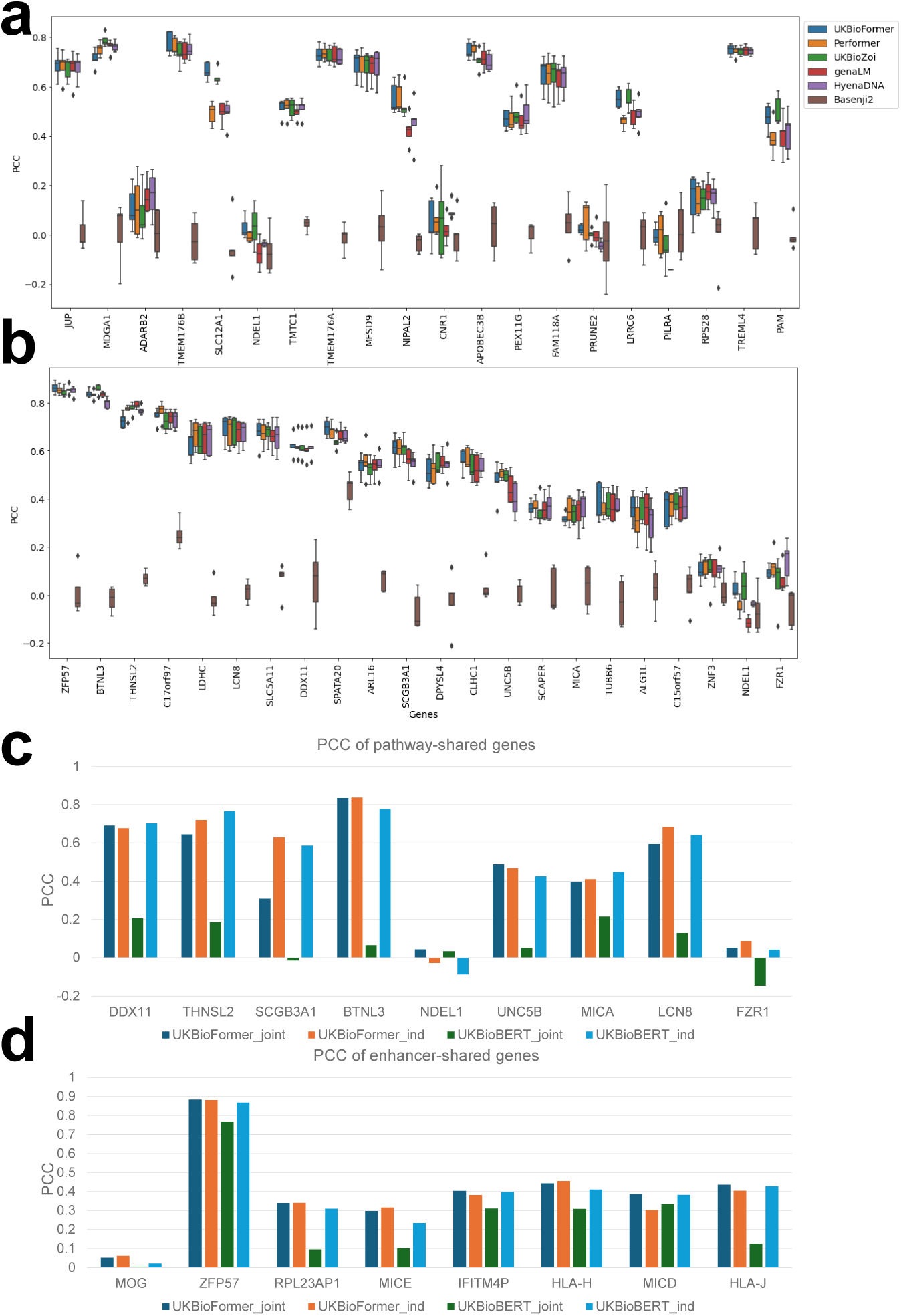
Performance comparisons among UKBioFormer and selected baselines. (a) Comparison of PCC across genes from group 1 for selected methods. (b) The comparison of PCC across genes from group 2 for selected methods. (c) Comparison between joint training (UKBioBERT joint, UKBio-Former joint) and independent training (UKBioBERT ind, UKBioFormer ind) for pathway-shared genes measured by PCC. (d) Comparison between joint training (UKBioBERT joint, UKBio-Former joint) and independent training (UKBioBERT ind, UKBioFormer ind) for enhancer-shared genes measured by PCC.

We also considered expression predictions across cohort-level data, for example, predicting the gene expression levels for African American (AA) individuals based on a model trained with European (E) individuals. According to Figure 4 (d) and Extended Data Figure 10, compared to training on Europeans and testing on Europeans, the prediction performance of UKBioBERT on the AA population was overall low, with prediction correlation negative for some genes. This may be due to the limitation of the embedding-based model. However, if we utilized Performer or UKBioFormer to make the same predictions, there was an obvious improvement in the prediction results based on the AA population, without modifying the training datasets. This result can be explained by the fact that sequence-based models can better capture variants that are correlated between groups while having a larger receptive field and therefore more generalization ability. Our analysis for the Religious Orders Study and rush Memory and Aging Project (ROSMAP) [65] dataset can be found in Appendix B.

It is well known that genes do not act alone, with genes of similar or complementary functions to regulate biological processes [66]. This is partly captured by gene-gene interaction [67]. Therefore, we hypothesize that training genes with similar functions together might boost each gene’s prediction performance. Furthermore, it has been demonstrated that joint training of different genes might improve prediction of some regulatory factors [7], and thus we explored if we can train UKBioFormer as a group. Therefore, we considered three different gene groups to examine this assumption, including the enhancer-shared group, the pathway-shared group, and the Enformer-pretrained group. The enhancer-shared group contains ZFP57 and other genes sharing the same enhancers in the blood tissue (n=8). The pathway-shared group contains genes in the same Gene Ontology (GO) pathway [68, 69] physiological response to stimulus (GO:0050896) (n=9). The last group contains genes used in the pre-training stage of Enformer (n=300). According to Figures 5 (c) and (d), the group-level training might not always benefit the expression prediction for genes in either enhancer-shared group or pathway-shared group, by comparing the group-level prediction performance with separate prediction performance using the same testing dataset. If we only use UKBioBERT, using a group of genes at the same time for training can even reduce the model’s performance. Moreover, according to Extended Data Figures 11 (a)-(c), the prediction performance based on the Enformer-pretrained group is also not promising, as many genes have a PCC score lower than 0.5. Therefore, the training model in the group-level genes for expression prediction is still challenging and could be affected by many limitations, for example, the low quality of training data as well as the underfiting of UKBioFormer. However, limited by the current data access and computation resources, these challenges are difficult to overcome. The results from Performer also suggest that the single-gene-level model can explain more variability than the group-gene-level model. Therefore, we believe that the current optimal plan is still training single-gene-level models for genes with good predictability.

Regarding the ablation studies and hyper-parameter tuning of UKBioFormer, we summarized the results based on the expression profiles of ZFP57 in Extended Data Figures 13 (a)-(d). In Extended Data Figure 13 (a), we demonstrated that the default setting referred from [7] to set epochs=100 is optimal, as increasing the epochs does not improve the model performance. We set the batch size to the maximum allowable value of the allocated GPU (8 for A40 and 12 for H100 or A100). Reducing the epoch is also not required and recommended as we have a validation dataset to select the model with the best weights. Extended Data Figure 13 (b) shows that using a lower learning rate can help improve the prediction performances. Extended Data Figure 13 (c) shows our results for the PEFT design. According to this figure, reducing the number of transformer layers can improve model performance, while other PEFT settings such as low-rank adaption (LoRA) [70] may reduce model performance. Furthermore, running PEFT design by reducing the number of transformer layers also outperformed the raw mode of Performer, and thus the optimal setting of PEFT design is to use the one-layer transformer architecture during the fine-tuning process. In Extended Data Figure 13 (d), we compared the default setting of the optimizer (Adam [71]) and loss function (mean squared error loss) with other ablation settings, by alternating optimizers (AdamW [72] or MUON [73]) or loss functions (pcc mse, huber loss [74], or group loss [7]), we did not see significant improvements, and thus the default setting of UKBioFormer is optimal. Using gradient accumulation (GA) to increase the batch size also does not improve model performance, shown in Extended Data Figure 13 (E). UKBioFormer also converged faster than Performer, shown in Extended Data Figure 13 (f). We finally compared UKBioFormer and UKBioZoi with implementation of Borzoi based on Flash Attention [75, 76], known as UKBioFlashZoi (the base model is FlashZoi [77]), and the results are shown in Extended Data Figure 14. According to this figure, UKBioFormer had the best performance, while UKBioZoi performed slightly better than UKBioFlashZoi.

### UKBioFormer can identify the direction of eQTL effects with the help of neural network explainability

Linking the variant with gene expression is a very challenging but important task, which can help us understand gene regulation. Previous research for eQTL inference requires access to matched genotype and expression data, which is expensive and time-consuming. Therefore, one of the major contributions introduced by the sequence-to-function model is to infer eQTL without having to measure hundreds to thousands of individual gene expression profiles, claimed by Enformer. However, the analysis of Performer shows that only using pre-trained Enformer cannot fully explain the effects of most eQTLs, and thus the fine-tuning process of the sequence-to-function model for better inferring eQTL is required and encouraged. Moreover, the fine-tuned model might also capture eQTL better than ElasticNet, especially for rare variants. Therefore, we hypothesize that UKBioFormer should also have the ability to identify eQTL with the correct direction and magnitude for the predictable genes. Furthermore, with the explainability tools designed for the sequence-to-function model, we can also compute gradients and perform In Silico Mutagenesis (ISM) to analyze the effects of different variants at the same locus. This approach us to uncover the role of such locus in regulating gene expression levels and to discover motifs which relates to gene regulation. The approaches we used to interpret UKBioFormer are summarized in Figure 6 (a).

**Fig. 6.**
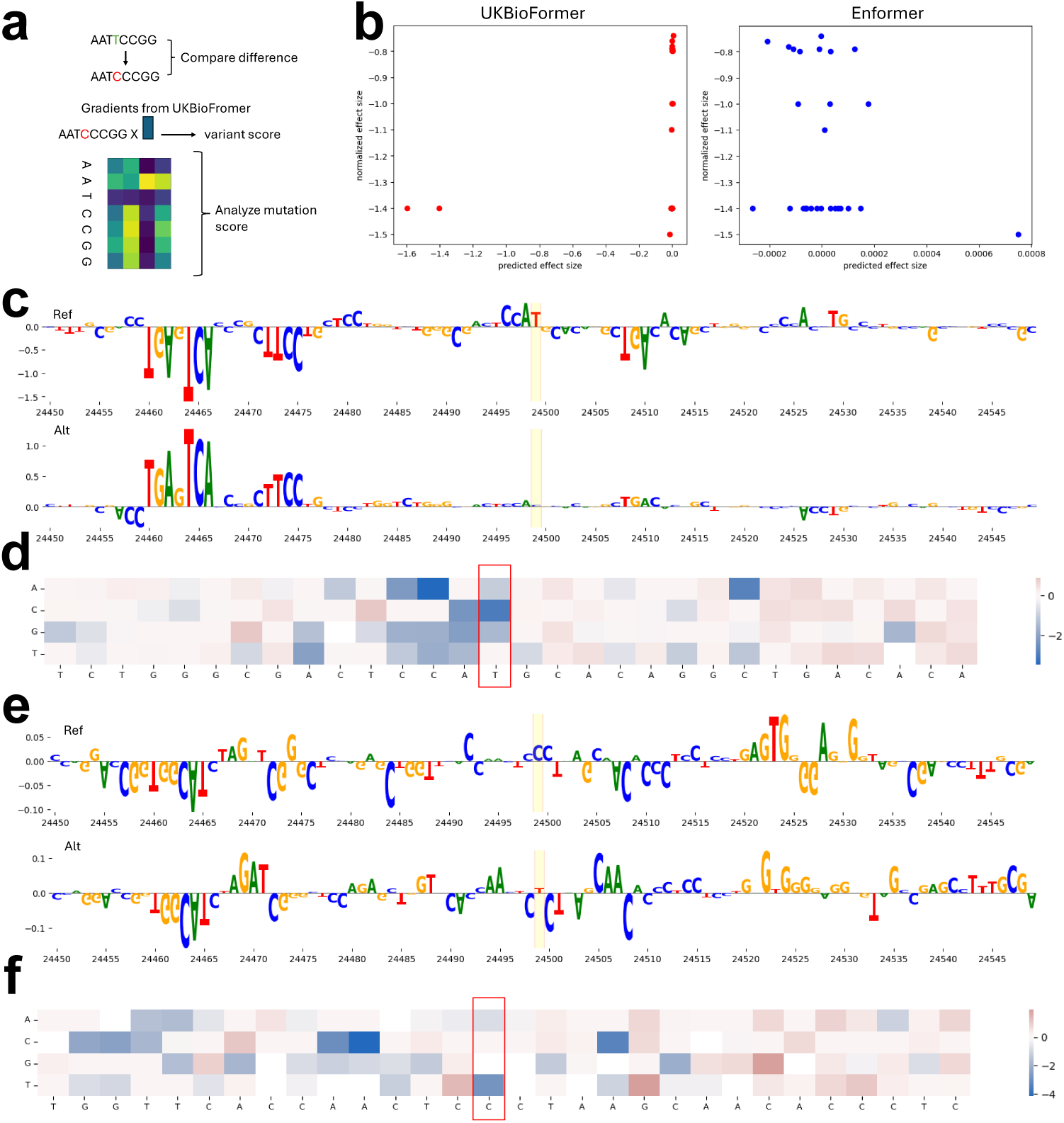
Understanding variant effects based on fine-tuned UKBioFormer. (a) The schematic diagram of the analysis of variance effects. We considered three approaches to analyzing variant effects, including effect size prediction, integrated gradients computation, and In Silico Mutagenesis (ISM). (b) The relationship between predicted effect size and normalized effect size based on the results from UKBioFormer and Enformer. (c) The visualization of base importance based on the reference sequence and the alternative sequence of eQTL with code rs9910080. (d) The results of ISM operation at the region near the location of rs9910080. (e) The visualization of base importance based on the reference sequence and the alternative sequence of eQTL with code rs9903086. (f) The results of ISM operation at the region near the location of rs9903086.

To identify eQTLs based on UKBioFormer, we performed a comprehensive benchmarking analysis between UKBioFormer and Performer. We also included AlphaGenome [78], which is a model with a much larger scale, as a baseline method of predicting eQTL effect. We first utilized the same training dataset from GTEx and selected the best model with the explainability mode based on the same validation dataset from blood. The estimated eQTL from the same dataset can be downloaded from the GTEx website, with information on reference alleles, alternative alleles, and normalized effect size. We then utilized ISM to predict the effect size for significant eQTL and compared the signs between the predicted and observed one. According to Extended Data Figure 15, UKBioFormer has a significantly higher proportion of eQTL with corrected signs for all 41 genes analyzed in this manuscript (p-value=0.02 between UKBioFormer and Performer, p-value=0.06 between UKBio-Former and AlphaGenome, one-sided Wilcoxon Rank-sum test). As a case study, we focused on the gene JUP as UKBioFormer has good predictions based on the testing datasets of JUP, and thus its expression levels are predictable. Firstly, we utilized the ISM method to predict the effect size of the top 30 eQTLs among 120 eQTLs with UKBioFormer ranked by p-value. Notably, 71% of predicted eQTLs have the correct signs, which is higher than the probability of random guess (50%) as well as the proportion estimated from Enformer (53%) or Performer (68%). Furthermore, we found that the prediction results of UKBioBERT for eQTL analysis also recovered the relationship between effect size and variance. We visualize the relationship between effect size and standard error (se) reported by GTEx database in Extended Data Figure 16, which shows a positive correlation between absolute effect size and se (PCC=0.49, p-value=1.72e-23). Such relationship was also reflected in the Extended Data Figure 9 from [79]. We observed that eQTL with high se computed by UKBioFormer are more likely to have correct directions (79% predicted eQTLs have correct directions for the high se group, and 64% predicted eQTLs have correct directions for the low se group), which matched the observed condition. We also plotted the relationship between predicted and observed effect size (normalized effect size) in Figure 6 (b) based on UKBioFormer (left panel) and Enformer (right panel), and UKBioFormer presented better alignment with observed effect size. We could also identify two representative eQTLs with perfect matching scores, including variant rs9910080 and variant rs9903086, which are also highlighted in the same figure. These two variants overlap with enhancer GH17J041791 of gene JUP, and thus, biologically they have the potential to affect gene expression levels. Second, we obtained the contribution of different base pairs to the prediction by running the gradient attribution method for a specific sequence region (inputs × gradients), the regions close to the TSS, and containing eQTLs of JUP. According to Figure 6 (c), we observed that the change of base pair for eQTL will cause an obvious gradient difference, and changing the selected base from a reference allele (T) to an alternative allele (C) results in a decrease in predicted gene expression, which matches the direction of this eQTL. We further considered running ISM to estimate the change of different alleles and their effects on gene expression levels, and Figure 6 (d) presents that the alternative allele also has the lowest ISM score. For variant rs9910080, we scanned the sequence with a motif database known as JASPAR [80] to discover associated motifs. By removing the motifs not in humans and ranking the discovered motifs with p-value, we find significant motifs such as JUN-class motif (TGAGTCAC, p-value=3.1e-5), which further support the capacity of UKBioFormer for selecting informative sequence patterns. Such conclusions can also be derived from our analysis of the other variant in Figures 6 (e) and (f). The motif overed based on the sequence close to variant rs9903086 is associated with zinc er factor (GCCGAGCCT, p-value=4.4e-4). The results of base importance and scores for selected variants, but computed based on Enformer and Performer are wn in Extended Data Figures 17 (a)-(d), which implies worse performance than BioFormer as the estimated variant effect is wrong in both magnitude and direct. Therefore, by using UKBioFormer to predict gene expression and capture the ifs associated with this expression phenotype, we can explore the effects of different mutations on gene expression and thus understand genetic relationships at a per level.

## Discussion

Sequence-to-function models have become important for understanding how genetic inforrmation can be executed or interpreted in various biological processes. This can help us further investigate the role of variants in expression regulation as well as gene heritability and predictability. However, few sequence-to-functions models focus on understanding individual genome information, and these models lack the design to capture the contributions of variants far from TSS as well as genome context. Furthermore, these models are large and inefficient, which creates challenges when deploying them in single-GPU platforms. Furthermore, incorporating genomic language models for interpreting variant effects at the individual level is also lacking in studies, even if this could be a promising direction as an enhancement of sequence-to-function models. Therefore, to overcome these challenges, we have proposed two sets of models to er understand functional genomics with the help of cohort-level genomic data. We rage UK BioBank database and extract the large-scale variants to train a gLM d on DNABERT2, known as UKBioBERT. We demonstrated that UKBioBERT generate a better representation for genes with functional annotations by bench-king the clustering performances with other gLMs. Such evaluation of function preservation in the latent space can also be used as a new metric for model pre-training. We also found that the embeddings from UKBioBERT are more informative to visualize individual information, enhance gene expression prediction in different cell s, and provide insights to gene predictability, which is not precisely explained by heritability. Therefore, variant information offers informative data augmentation for training.

To link UKBioBERT with sequence-to-function models, we fused the embeddings UKBioBERT with models including Enformer or Borzoi and proposed two new sequence-to-function models, known as UKBioFormer and UKBioZoi, to predict gene ression for different individuals more efficiently. We demonstrated that these two models had good performance in the genes with high predictability, and their results had high correlations with ElasticNet and outperformed Performer. Based on results, we recommend using UKBioZoi when computation resource is a challenge, while UKBioFormer is expected to achieve a better performance. Moreover, we wed that using UKBioFormer can also make cross-cohort expression prediction, example, training a model from individuals with European ancestry and making prediction in African-American individuals, which demonstrates that UKBioFormer is less affected by the biases brought by different variants across cohorts. We also discussed in detail the design options for UKBioFormer, including how to choose the loss functions, optimizers, hyper-parameters, and plans of PEFT. Therefore, work may lead to future development to combine these two types of models to make further improvements or more interesting applications.

Finally, the explainability of UKBioFormer can also explain the role of variants in regulating gene expression levels with good predictability. We demonstrated that UKBioFormer has the ability to predict the correct direction of over 70% eQTLs, and can also match the observed effect size well for certain important eQTLs. Furthermore, we can also utilize UKBioFormer to test the effect of base changes on the role of surrounding bases in gene expression regulation and select target variants to test their effects experimentally. Therefore, we believe that UKBioFormer will open more possibilities to explore how variants affect gene expressions.

For future work, we plan to tackle one challenge discussed in this study, namely, the challenge of effective multi-gene training for cross-individual prediction. We found that training the model with multiple genes cannot always improve expression prediction for all the selected genes in different scenarios. This could be a problem due to the drawbacks in model design, the limitation of computation resources, or the noise in data. As a next step, we plan to focus on cell-type information across individuals and incorporate more modalities as training datasets to see if we can overcome this challenge in the cell-type-specific context. Moreover, it is also promising to link the discoveries of UKBioBERT and UKBioFormer with therapeutic research in diseased samples, which might inspire novel treatment plans for diseases associated with genetics.

## 4 Methods

### Problem statement

Here we intend to first have a model M, which can model the DNA sequence dataset D by learning a good representation of arbitrary sequences. This approach can be implemented by the self-supervised approach (e.g., mask language modeling strategy [16]) or supervised approach (e.g., predicting Linkage Disequilibrium (LD) score [33]) at the pre-training stage. Furthermore, we also intend to have the other model P, which is built based on the pre-trained model M and should predict the gene expression levels based on the corresponding sequence information of the given gene in the population level. To achieve this goal, we can train the model P by minimizing the distance between the observed gene expression levels *y* and predicted gene expression levels 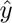.

### Implementation of UKBioBERT

Here we discuss the model architecture and pre-training strategies for UKBioBERT. We utilize DNABERT2 [17] (which is a Bidirectional Encoder Representation from Transformers [16] model pre-trained with reference genome), and continue pre-training our method based on the sequences with variants. We create a function to modify the reference genome with matched variant information, as shown in Algorithm 1. The variants come from UK BioBank [33] database’s EUR population. Our pre-training strategy is masked language modeling, which is determined after comparing it with the contrastive-learning-based training approach and predicting the LD score approach. We split the training, dation, and testing dataset for UKBioBERT to select the optimal checkpoint, and test the clustering score. The loss function for model training is to classify the k token by minimizing the cross-entropy loss.

**Algorithm 1.**
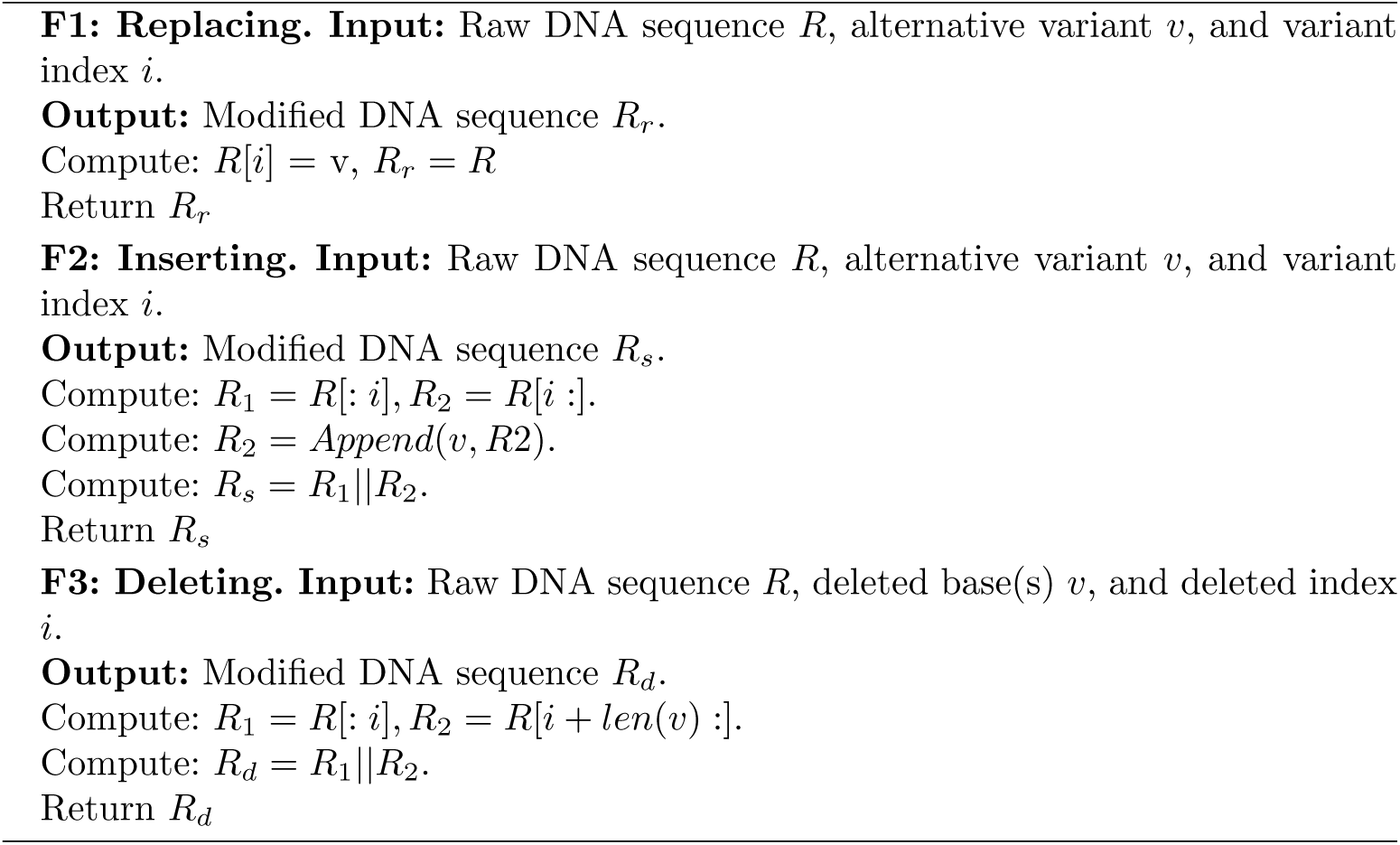
Sequence editing functions.

We follow the default setting of DNABERT2 for hyper-parameter selection. We ect 13 million variants and paired sequences to construct the sequence dataset. en performing the pre-training experiment, we randomly split the collected ence dataset into training/validation/testing sets with a proportion 0.8/0.1/0.1. BioBERT supports sequences with different lengths, as long as the user has paired U capacity. Based on our experiments, an H100 GPU can support one 100K DNA ence as input. We also pre-train HyenaDNA [2] with a causal language modeling roach based on sequence information from UK BioBank data as a baseline, which enoted as UKBioHyenaDNA.

### Applications of UKBioBERT

We extract the embeddings of DNA sequences UKBioBERT for downstream applications.

1. Visualization. We utilize the DNA sequence embeddings corresponding to gene tional labels or individual labels for visualizing the DNA sequence in the lower ension, which shows reasonable clustering performances.
2. Cell-state-specific expression prediction. UKBioBERT can generate sequence eddings to help EPInformer [15] in predicting gene expression levels for known cell s, including K562 and GM12878 [81]. These cell lines contain accessible information nhancers, chromatin interactions, etc. We encode one promoter sequence and 60 ancer sequences with length 2000 (the default setting of EPInformer) and merge the embeddings with the default input of EPInformer to provide extra information for model training. Our embeddings can significantly improve model performance, compared with the default mode.
3. Individual gene expression prediction. Leveraging gene embeddings from UKBioBERT, we can also use the encoded gene sequence from Whole Genome Sequencing data [82] of individuals to predict the corresponding expression levels in the selected sample for the corresponding gene. Our regressor is ElasticNet [83] and we perform cross-validation to select the best hyper-parameter setting and make predictions for five different testing sets.
4. Individual stratification. The gene embeddings from UKBioBERT can further help us understand the differences across individuals and how they affect the prediction performances. We consider different labels, including expression levels, ancestries, and genders to explore the possibility of using UKBioBERT to distinguish them. Our results show that UKBioBERT can link the predictability of gene expression levels with the embedding quality, which further helps us to determine which set of genes can be successfully predicted.

Finally, we intend to highlight that UKBioBERT is powerful in generating sequence embeddings, and thus its application is not limited to expression-related analysis.

### Implementation of UKBioFormer

We build a new transformer-based model, named as UKBioFormer, based on Enformer [30] and UKBioBERT. The improvement of UKBioFormer is discussed in two sections, the design contribution as well as the engineering contribution. UKBioZoi follows a similar idea but its base model is Borzoi, so we do not plan to discuss it in detail.

Regarding the design contribution, previous research demonstrated that the attention mechanism applies a higher weight to the transcription starting site (TSS) and to regions in close proximity to the TSS, which might not model correctly for the contribution of far-away genetic information or variants [7, 84]. Therefore, incorporating the gene embeddings from UKBioBERT can further help the model to capture the nucleotide information in marginal regions, as we encode DNA sequences by averaging the embeddings of different tokens and our pre-training strategy does not include the TSS information to reduce inductive bias. In summary, we believe that UKBioFormer can make a good balance between learning the local and global genetic information for expression prediction.

Regarding the engineering contribution, we consider pre-training, parameter efficient fine-tuning (PEFT) [85], and feature fusion to enhance the efficiency and capacity of UKBioBERT. First, we do not train UKBioFormer from scratch but utilize the model weights from Enformer as initial parameters. According to [7], this approach might at least help in predicting the expression levels from genes pre-trained based on the CAGE-seq of Enformer, and thus starting from a good initial weight can also help the model perform better transfer learning. Second, we consider applying the PEFT approach for fine-tuning UKBioFormer with lower resource requirements as well as possibly better model performances. Using PEFT can also support the model with longer sequences as inputs, compared with the 49k length used by Performer for fine-tuning, our UKBioFormer can accept over 100k length sequences. In addition, we e tried the low-rank adaption (LoRA) [70] approach and pruning approach to optimize the parameter usage of UKBioFormer, and found that pruning the transformer rs can improve the training efficiency as well as performances better than LoRA ur Results section. Compared with UKBioBERT, UKBioFormer also has a faster vergence speed, as discussed in the Results section. Third, we demonstrated that on is an effective approach to combine the gene embeddings from UKBioBERT our learning target. Here we finally extract the outputs from the regression head of embeddings from pruned Enformer (which is training) and weighted average the puts from such gene embeddings and the outputs from the regression head based he gene embeddings from UKBioBERT (which is frozen). The weight is learnable always in the range of (0,1). This approach can not only guarantee that our modwill capture the local and global genetic information but also will not affect the ult abilities of interpreting UKBioFormer.

The loss function of the fine-tuning method is mean squared error loss, which etermined after comparing different kinds of loss functions. Here we evaluated functions including mean squared error (MSE) loss, the joint loss between MSE PCC (pcc mse), huber loss (huber loss) [74], and group loss (group loss) [7]. To truct pcc mse, we train the model by maximizing the correlation between the rved and predicted gene expression levels in the training dataset while minimizing MSE between them. To construct huber loss, we minimize the balanced MSE loss ween the observed and predicted gene expression levels. To construct group loss, not only minimize the MSE between the observed and predicted gene expression, minimize the MSE between the difference of individuals in two groups to let the el also learn the variability existing in individuals.

Most of our hyper-parameters follow the settings from [7]. We have a smaller h size because we do not have the computation resources used in this manuscript H100). However, our model’s performance for cross-individual prediction is comble and even better with their setting. Our method is implemented based on LU [86]. The parameter size of UKBioFormer is 230.7 million. The parameter size kBioZoi is 170.7 million.

### Applications of UKBioFormer

Here we discuss the application scenarios of BioFormer, which cover the important tasks in understanding functional genomics, ing from expression prediction to expression quantitative trait locus (eQTL) identification.

1. Cross-individual gene expression prediction. By fine-tuning UKBioFormer with ning individuals and validation individuals, we can obtain an optimized model specific genes’ expression prediction. We can further test our model’s perforce based on the DNA sequences from the testing individuals. We performed fold cross-validation setting to report the variance, that is, we split training/validation/testing with the ratio 8:1:1 for each fold, and report the metrics based on ing data. We demonstrated that our methods can outperform baseless ones for icting the gene expressions with good predictability.
2. Reducing biases in expression prediction. We also demonstrate that our methods improve prediction across individuals with different confounder variables, such as estry. We train our model with individuals from European (E) ancestry and test the model with individuals from African American (AA) ancestry. It shows that our fine-tuning approach achieves better performance for certain genes, compared with selected baselines. We also notice that for this application, using all of the transformer layers provided a more stable and better prediction on average.
3. Testing the synergetic effect of the training model at a group level. We also explore the potential contributions brought by training genes with similar features in a group and investigate if this approach can enhance the prediction performances for all selected genes or not. Here we consider three different groups, including enhancer-shared group, pathway-shared group, and Enformer-pretrained group. The first group contains genes sharing the same enhancer in the blood tissue, the second group contains genes in the same Gene Ontology (GO) pathway, and the last group contains genes pre-trained in the Enformer model, provided by [7, 30]. We compare the joint-training results with individual-training results to analyze the prediction results across different genes, and make conclusions based on our prediction results accordingly.
4. Interpreting mutation effects. We can also utilize tools to interpret the contributions of mutation to the change of gene expression prediction. Our mode accepts frozen gene embeddings paired with DNA sequence information as inputs, and thus the explainability analysis can be conducted with DNA sequences and the sequence encoder. Here we compute the gradient x inputs [30], which is a type of attribution, for each base to explain the base-level contribution for gene expression prediction. The gradient is computed based on the absolute value of the gradient of predicted expression levels. We also collect the reference sequences and alternative sequences and utilize In Silico Mutagenesis (ISM) [30, 86] to compute the weights for different sequences. By comparing the vector of weeds for these two cases, we can understand whether the given variant or mutation can enhance or reduce gene expression levels.
5. Identifying eQTL. The traditional approach utilizes a statistical framework to estimate the contribution of a single variant for affecting gene expression prediction, known as eQTL inference. Different eQTLs might have different directions and magnitudes for the targeted gene. We demonstrated that our methods can help predict the directions of eQTLs for genes with good predictability, and improve the prediction of magnitudes for certain eQTLs with strong effect size (which is defined as a large absolute value). However, there are still various challenges for further identifying reliable eQTLs.

### Metrics

For the model pre-training stage, we evaluate the quality of embeddings based on the preservation of the gene function context. The idea is to run a clustering algorithm based on gene embeddings from different foundation models and compare the cluster labels with annotated gene functional labels. We access the optimal resolution of the Leiden clustering algorithm for each method’s generation to perform a fair comparison. The metrics we considered included Normalized Mutual Information, Adjusted Rand Index, Average Silhouette Width, and their average score Avg. These metrics are computed based on scikit-learn [40] and scIB [41]. Details are discussed below.

- Normalized Mutual Information (NMI): NMI is a score to evaluate the performance of biological information conservation. We compute this score based on the mutual formation between the optimal Leiden clusters and the known cell-type labels and then take the normalization.
- Adjusted Rand Index (ARI): ARI is a score to evaluate the performance of biological formation conservation. ARI is used to evaluate the agreement between optimal eiden clusters and gene labels.
- Average Silhouette Width (ASW): We have gene type ASW for this metric. For one cell, ASW calculates the ratio between the inner cluster distance and the intra uster distance for this gene. Therefore, higher *ASW_cell_* means better biological formation conservation. To make them consistent, we take the normalization, that

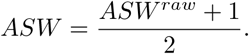

All of the selected metrics in this section range from 0 to 1, and thus we can also age them to evaluate the effect jointly, which can be represented as Avg ∈ (0, 1). For the cell-type-specific gene expression task, we utilize the Pearson Correlation Coefficients (PCCs) between the measured gene expression levels and observed gene ression levels as the metric, which follows the default choice of EPInformer.

For the individual gene expression task, we utilize PCCs, *R*^2^ score, and mean square error (MSE) as metrics for evaluation. Higher PCCs and *R*^2^ scores represent a better el, while higher MSE represents a worse model. All the metrics are implemented d on scikit-learn or Scipy [87].

For the eQTL discovery task, we take the observed eQTL information from GTEx site [82], including the variant information and normalized effect size (NES). We pare the signs and absolute values of NES and predicted effect size to perform the parison.

### Baselines

Here we discuss the baselines used in this study for different tasks. For the model pre-training stage, we consider various foundation models, which are ned based on different strategies as well as base models. The first group is multi-biological sequence foundation models, including DNABERT [11], DNABERT2 GPN [28], HyenaDNA [2], nucleotide transformer (NT) [42], Grover [43], a LM [19], Caduceus [44], ESM [45], METAGENE-1 [46], LucaOne [47], and Evo2 Most of the models are pre-trained with DNA sequences, while LucaOne takes ti-omic-sequence information in the pre-training stage and ESM utilizes the gene’s esponding protein sequence as inputs. If they have multiple versions, we utilize model with the largest parameter size. The embeddings from ESM are extracted d on the file provided by KGWAS [88]. These models generate sequence embeds or novel DNA sequences, which can be used for further applications. The second p is an expression-based large-scale sequence model, including Enformer [30] and zoi [48]. These models are trained to predict gene expression levels under differtechnologies (CAGE-seq or RNA-seq), and the learned embeddings will encode functional information of the selected sequences. The embeddings and outputs of e models are often used for tasks related to gene expression prediction. The last p is the general foundation model, for example, Llama 3.1 [49, 50] was pre-trained multi-language data, which is a powerful open-source tool for tasks related to ural language processing. Since Llama 3.1 does not provide direct access for its generated embeddings, we pre-train Llama 3.1 with DNA sequence information from UK BioBank and use AngIE [89] to generate the embeddings from Llama 3.1.

For the cell-type-specific gene expression prediction, we consider EPInformer [15] and its variants as baselines. EPInformer is a transformer-based predictor leveraging the information of promoter sequences, enhancer sequences, as well as epigenomic information for enhancing model performance in gene expression, enhancer identification, and variant effect estimation based on data from cell lines. In addition to our final design, we compared EPInformer which incorporates different features, including the combination of EPInformer and HyenaDNA, scELMo [57], and EpiBERT [56].

For individual gene expression prediction, we include ElasticNet, UKBioBERT, Gena LM, HyenaDNA, and Basenji2 as baselines. ElasticNet is widely used in predicting gene expression at the individual levels based on SNPs. We also utilize UKBioBERT to generate sequence embeddings and these embeddings can be treated as inputs of ElasticNet to make predictions. We extract gene embeddings based on Gena LM and HyenaDNA and train with ElasticNet for expression prediction. We use the zero-shot mode of Basenji2 with pre-trained weights to infer gene expression profiles from sequences. We use the cross-validation setting of ElasticNet and 2000 maximal epochs for model training [7], and we only consider single-gene model for both ElasticNet and SuSiE [90] in our experiment. We also consider the variants of Borzoi as UKBioZoi and include them in our benchmarking analysis.

Details of the parameter setting of baseline models are summarized in our codes.

### Datasets

The dataset used in pre-training can be downloaded from the UK BioBank website. The gene expression and sequence information of cell lines K562 and GM12878 can be downloaded from the links provided by EPInformer. The dataset used in individual gene expression prediction can be downloaded from GTEx and ROSMAP websites. Access to datasets from UK BioBank, GTEx, and ROSMAP requires extra requests. We summarized the dataset information in Supplementary File 1.

### Code availability and reproductivity

For pre-training data pre-processing, we reply on Yale High-performance Computing Center (YCRC) and utilize one CPU with up to 800 GB. For model pre-training, we utilize one NVIDIA A100 (H100) GPU with up to 100 GB RAM. For model fine-tuning, we utilize one NVIDIA A100 (H100/A40) GPU with up to 70 GB RAM.

The codes of this project can be found in https://github.com/HelloWorldLTY/UKBioLM. The license is MIT license.

## 5 Ethics and Inclusion

Although UKBioBERT and UKBioFormer are not biased to gender, races, and other factors, the users are solely responsible for the content they generate with models in UKBioBERT and UKBioFormer, and there are no mechanisms in place for addressing harmful, unfaithful, biased, and toxic content disclosure. Any modifications of the models should be released under different version numbers to keep track of the original models related to this manuscript.

The target of current UKBioBERT and UKBioFormer only serves for academic arch. The users cannot use it for other purposes. Finally, we are not responsible any effects of the use of the model.

## Acknowledgments

We thank Shiron Drusinsky and Sean Whalen for the suggestions of data preprocessing and the reproduction of Performer and ElasticNet, and Dr. Zihan Dong and Leqi Xu for the suggestions of using UK BioBank data. We thank NSF ACCESS for providing computation resources to support our research. This research has been conducted using the UK BioBank Resource under Application Number 29900. This project is supported in part by NIH grants U24HG012108 and U01HG013840.

## Author contributions

T.L. proposed the study. T.L. and X.Z. ran all the experiments. R.Y. and H.Z. provided the computation resources. J.L. and L.P. provided cellline datasets. All authors wrote the manuscript. H.Z. supervised this project.

## A Analysis of genome-level factors to explain gene predictability

In this section, we discuss our process to explore the gene predictability from a deeper and more biology-centered approach, that is, linking the observed predictability with genome-level information. We selected several potential factors which might affect the predictability of genes, including location of genes in the chromosomes (GL), number of unique sequences (NUS), number of enhancers/promoters (NEP), and GeneCards Inferred Functionality Scores (GIFts) [62], as our extra experiments. The computation of GIFts is based on the number of functions of the given gene divided by the total number of gene functions. The predictability is defined by the PCCs of selected genes used in our results section. To test the effect of gene location information, we separated the genes into odd group (location is in odd-index chromosome, e.g., chr1) and even group (location is in even-index chromosome, e.g., chr2) and performed Mann–Whitney U test, and the boxplot for the PCCs of these two groups is shown in Extended Data Figures 12 (a) and (b). For other factors, we show the scatter plot in Extended Data Figures 12 (c)-(e), and we also computed the PCC and SCC between gene predictability and these factors. Based on our results, we found that only GIFts showed a significant correlation with gene predictability. The gene predictability did not show significant difference in different location groups, as well as strong correlation with other factors. Therefore, the complexity of gene functions can negatively affect the gene predictability.

## B Analysis of testing gene expression profiles from ROSMAP dataset

In this section, we test the cross-cohort prediction for ROSMAP brain cohort RNA array data by training Performer based on GTEx brain cohort RNA-seq and paired WGS data, and our genes used for testing are provided by the original manuscript of Performer. As we observed strong prediction correlation between Performer and UKBioFormer, we believe that the training results of Performer can be a good indicator to determine if there exists reward based on the joint training current setting. We visualize our training results in Extended Data Figures 11 (d)-(f), and the statistics show that the prediction performances are not good enough, as many genes’ PCC scores are lower than 0.5. We also found similar conclusions from the testing results based on ElasticNet or SuSiE [90] (Extended Data Figures 11 (g) and (h)), while Performer has a higher average PCC. The low R2 and high MSE can be explained by magnitude difference caused by sequencing technologies. Furthermore, one possible reason for explaining both case is the low variance of testing data, as we found that in the case of across-individual prediction, the largest variance of the tested genes was 0.29 and the smallest was 6e-4. Since the vast majority of the gene expression levels were between 7 and 8, it can be assumed that there is little difference across individuals. This discovery demonstrated that the cross-cohort prediction under the multiple-gene setting is still very challenging.

## C. Supplementary Figures

**Extended Data Fig. 1.**
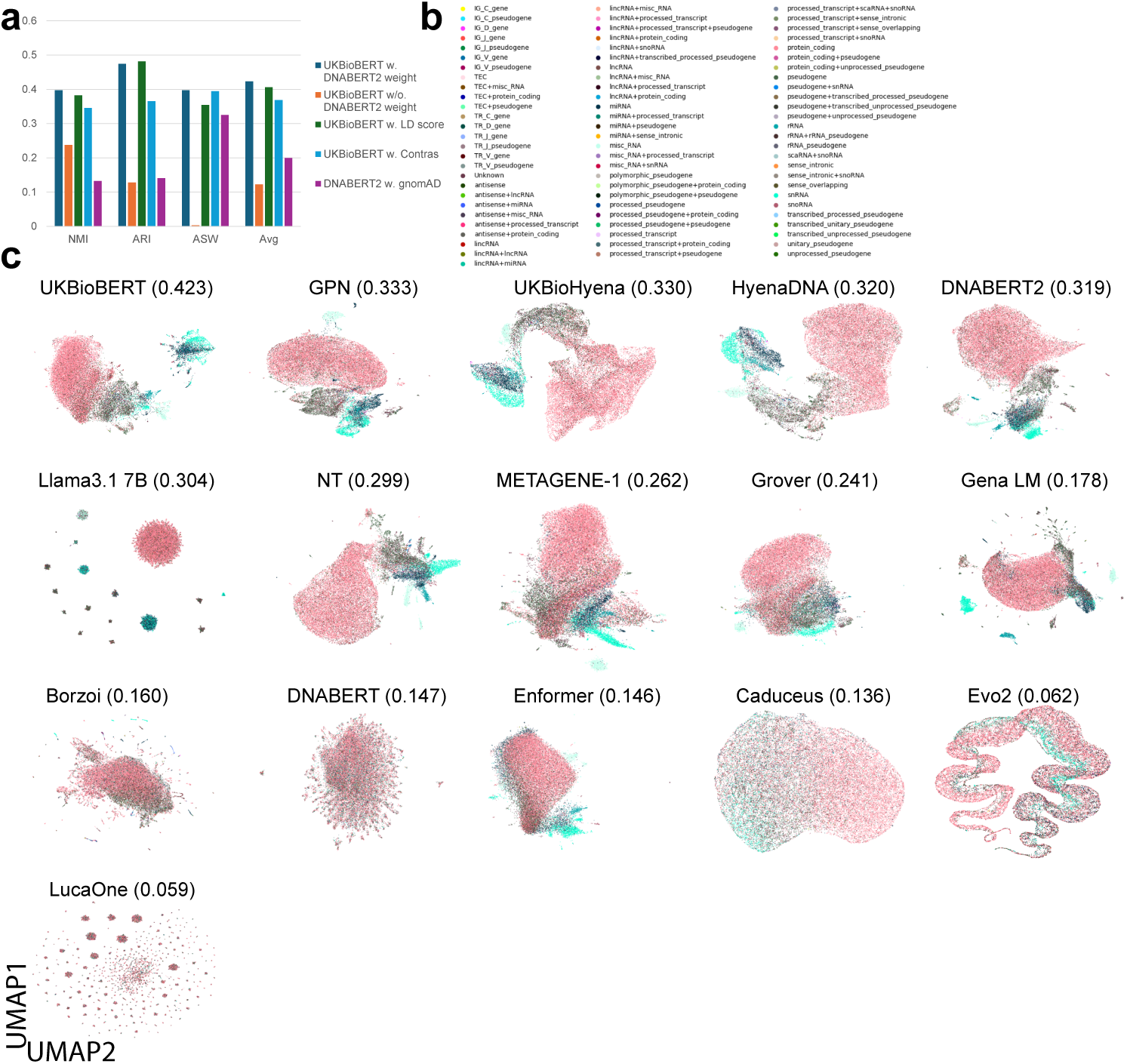
Comparisons of pre-training strategies and visualizations of baselines. (a) Clustering scores of UKBioBERT based on different pre-training strategies. (b) Gene function labels used in this manuscript. (c) UMAP visualizations of the full list of baselines.

**Extended Data Fig. 2.**
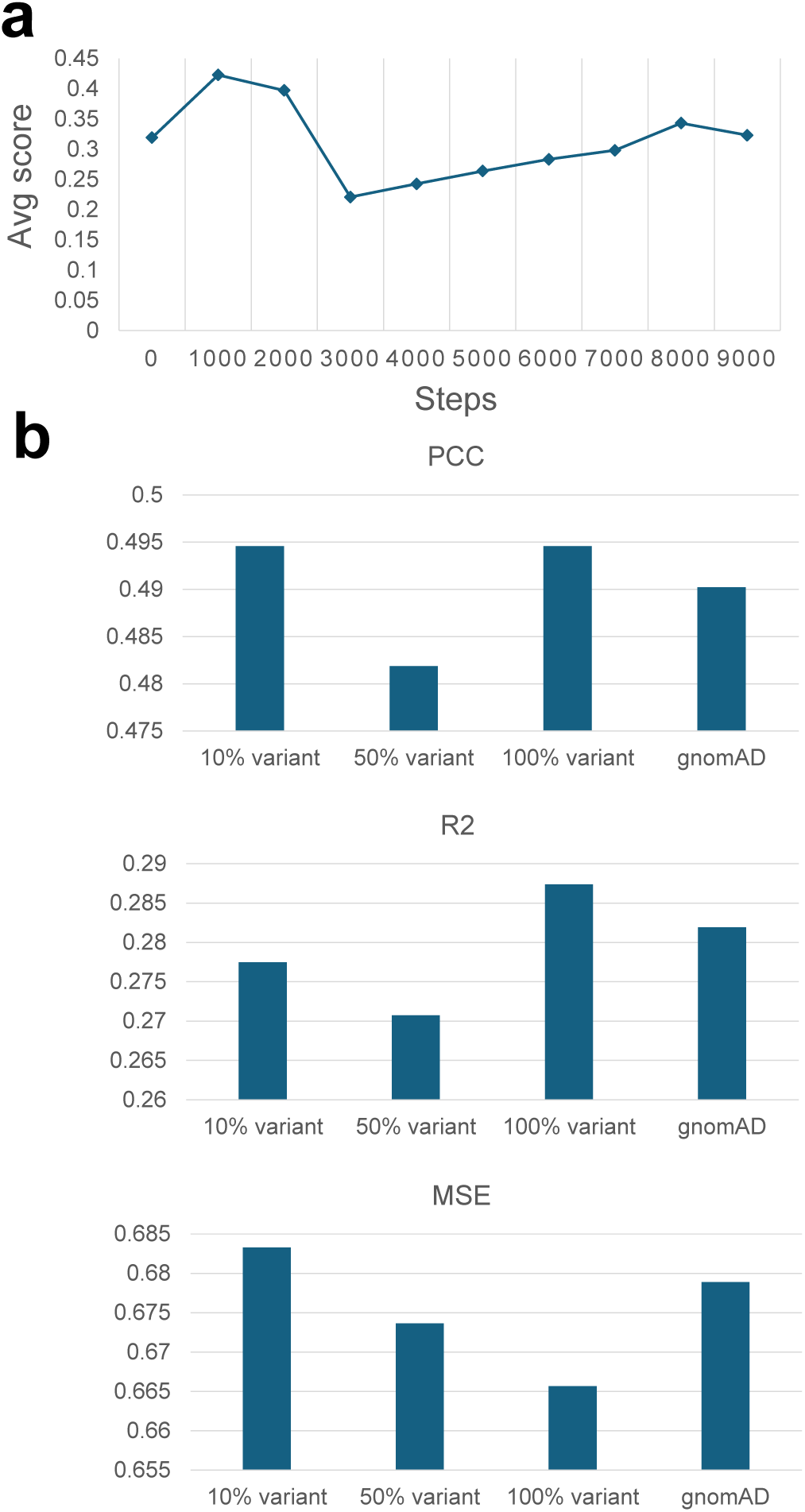
Details of pre-training experiments. (a) The score vs. training step curve for UKBioBERT. (b) Data scale comparison based on the gene expression level prediction task.

**Extended Data Fig. 3.**
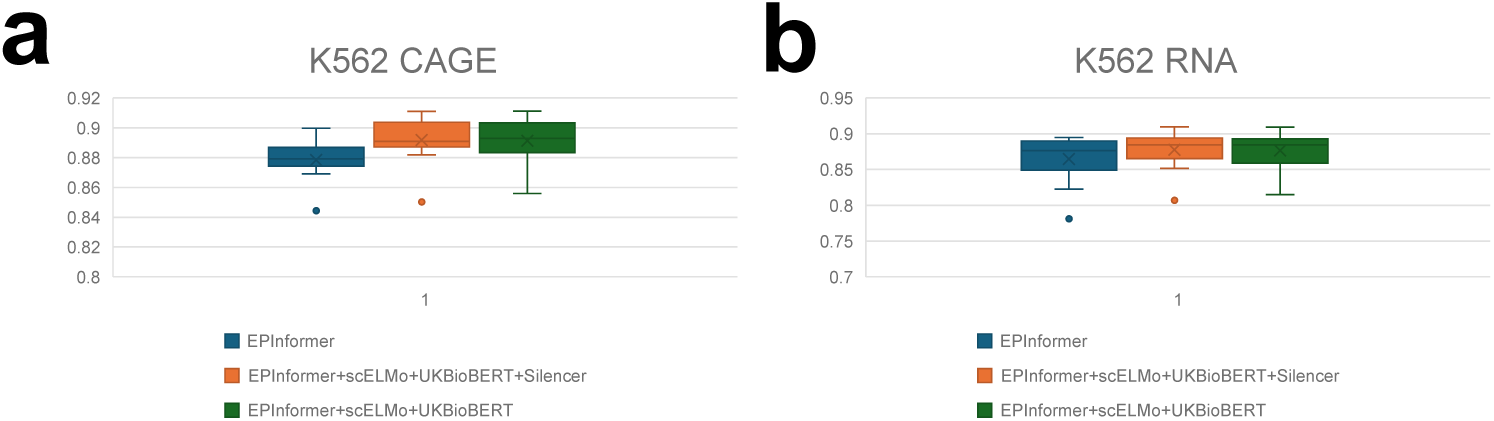
Comparisons of modified EPInformer with different features including silencers. (a) Prediction performances of different methods based on K562 cell lines from CAGE-seq. (b) Prediction performances of different methods based on K562 cell lines from RNA-seq.

**Extended Data Fig. 4.**
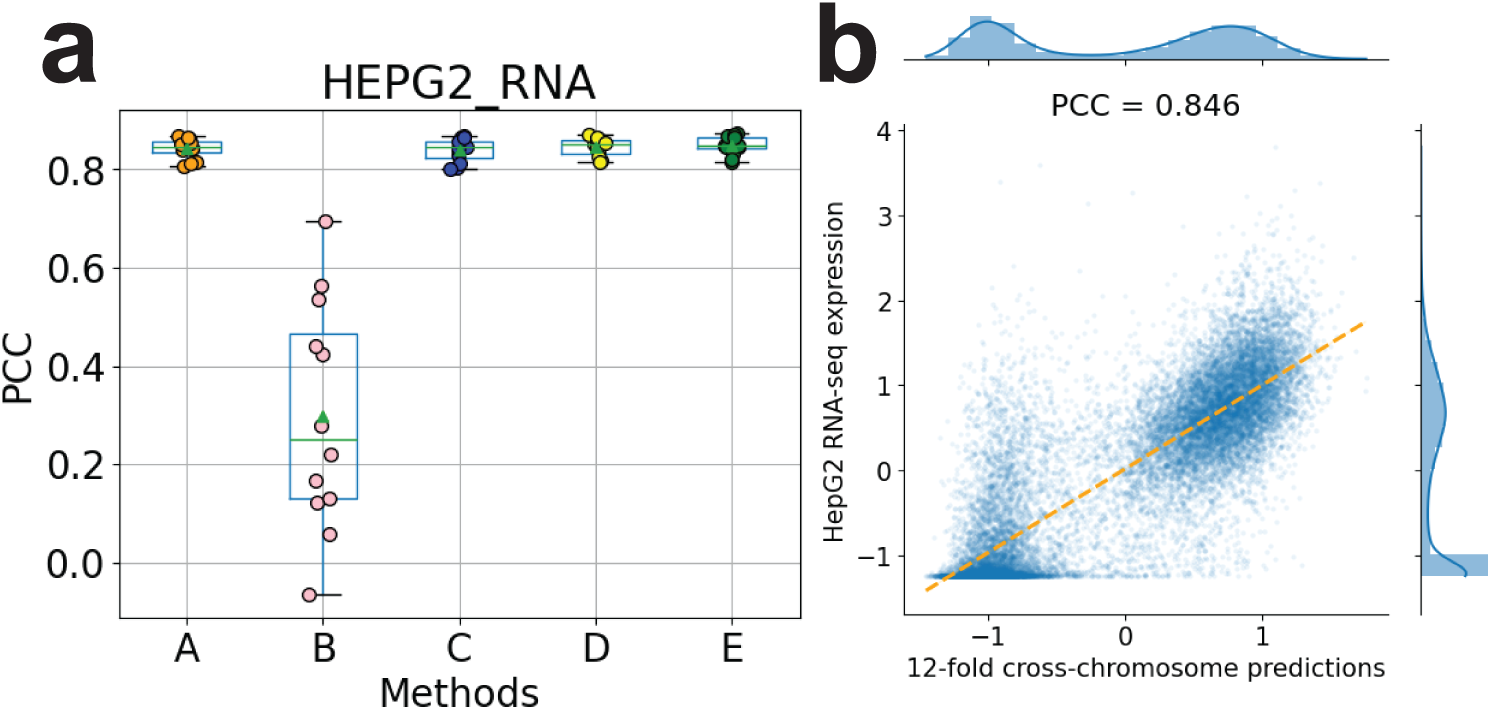
Comparisons of modified EPInformer with different features for predicting gene expression levels in HepG2 cell line. (a) Prediction performances of different methods based on HepG2 cell line from RNA-seq. The mode A represents default EPInformer, the mode B represents EPInformer+EpiBERT, the mode C represents EPInformer+UKBioBERT, the mode D represents EPInformer+scELMo, and the mode E represents EPInformer+UKBioBERT+scELMo. (b) Relationship between the predicted and observed expression levels based on EPInformer with gene embeddings from UKBioBERT and scELMo.

**Extended Data Fig. 5.**
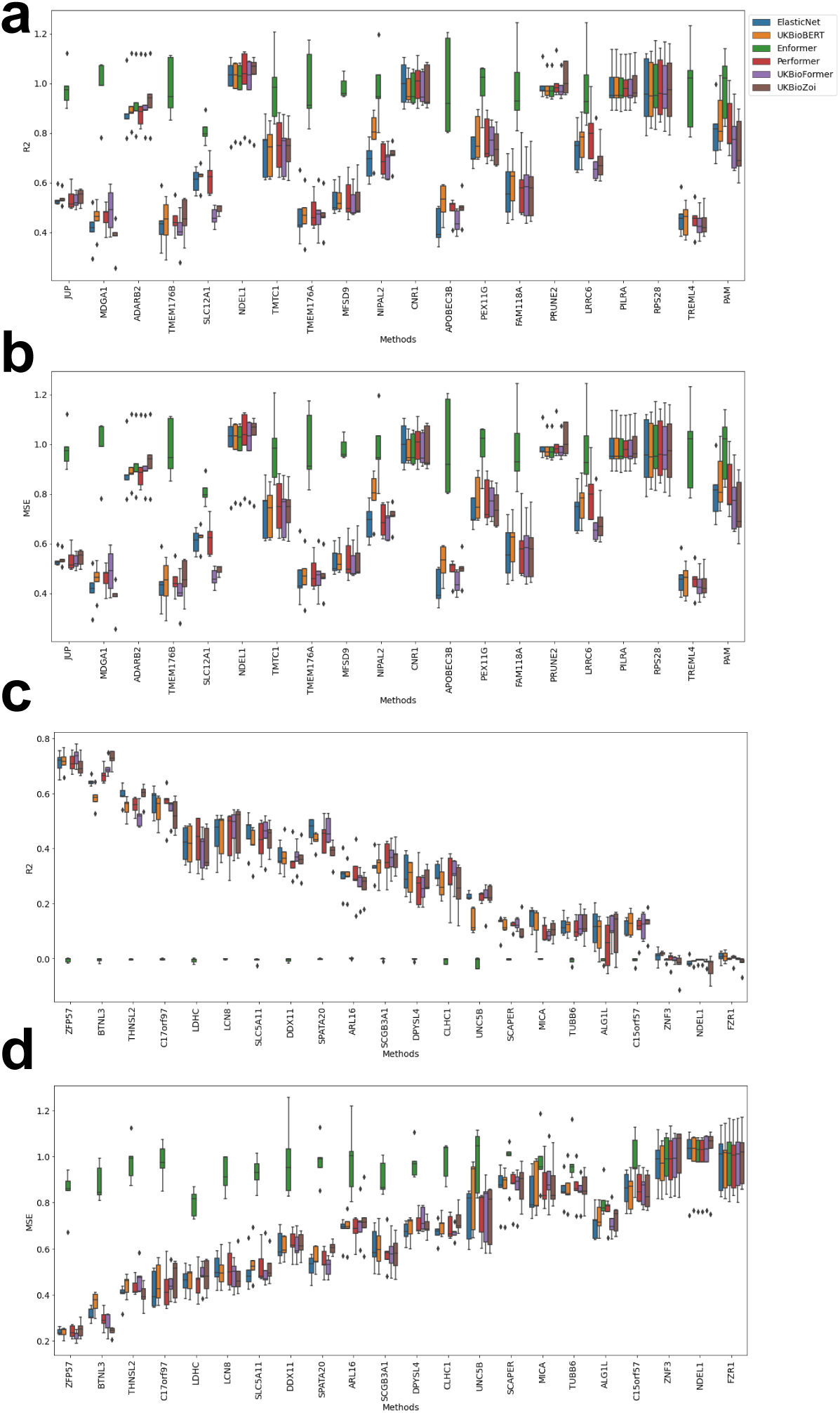
Comparisons of different methods across selected genes measured by R2 and MSE. (a) R2 scores of different methods across genes in group 1. (b) MSE scores of different methods across genes in group 1. (c) R2 scores of different methods across genes in group 2. (d) MSE scores of different methods across genes in group 1.

**Extended Data Fig. 6.**
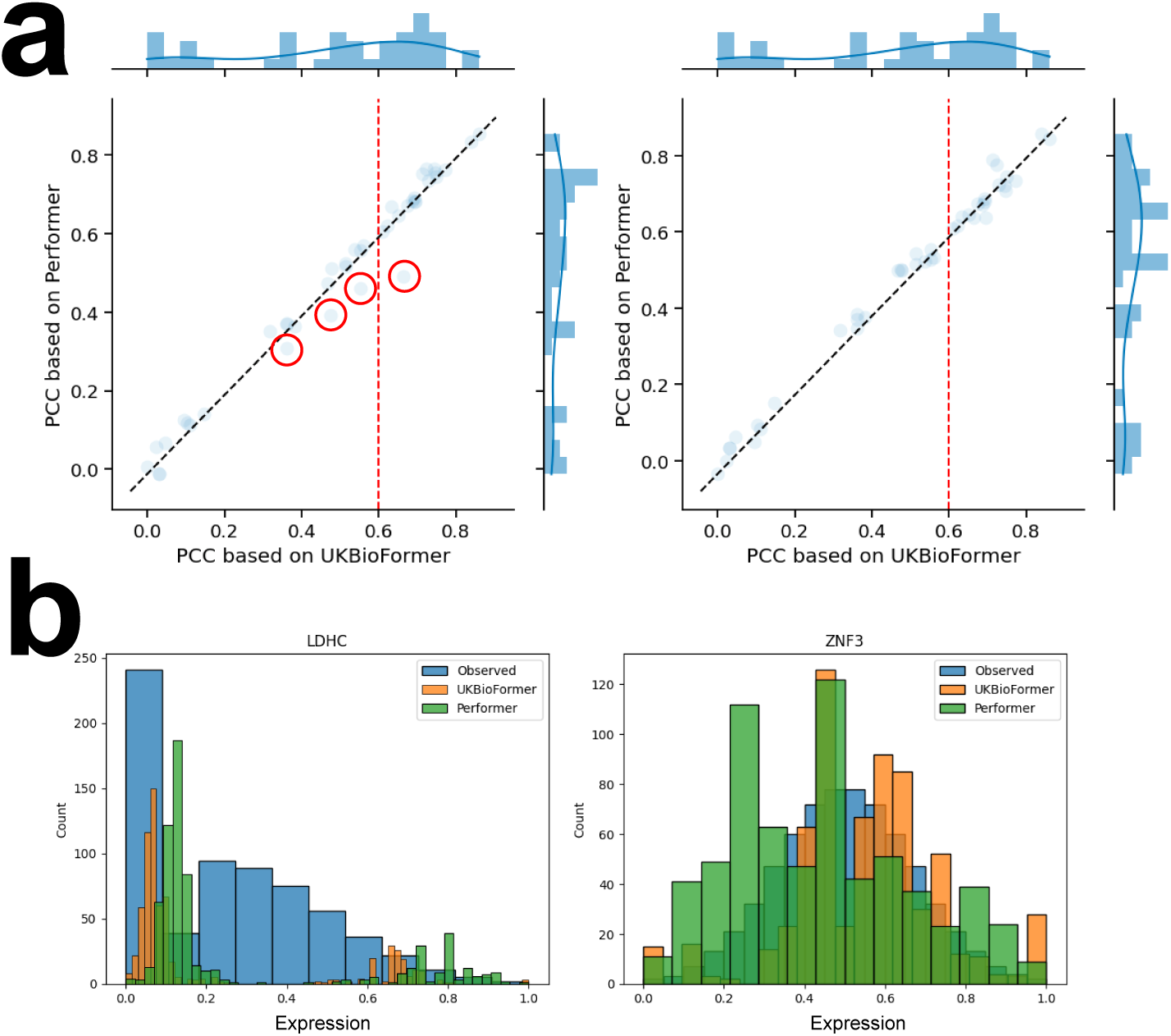
Comparison of paired methods for gene expression prediction. (a) In the scatter plot, each dot represents one gene. The left panel represents the comparison of PCCs between UKBioFormer and Performer. The right panels represents the comparison of PCCs between UKBioFormer and UKBioZoi. (b) The detailed expression comparison based on the distributions between UKBioFormer and Performer for the highlighted genes.

**Extended Data Fig. 7.**
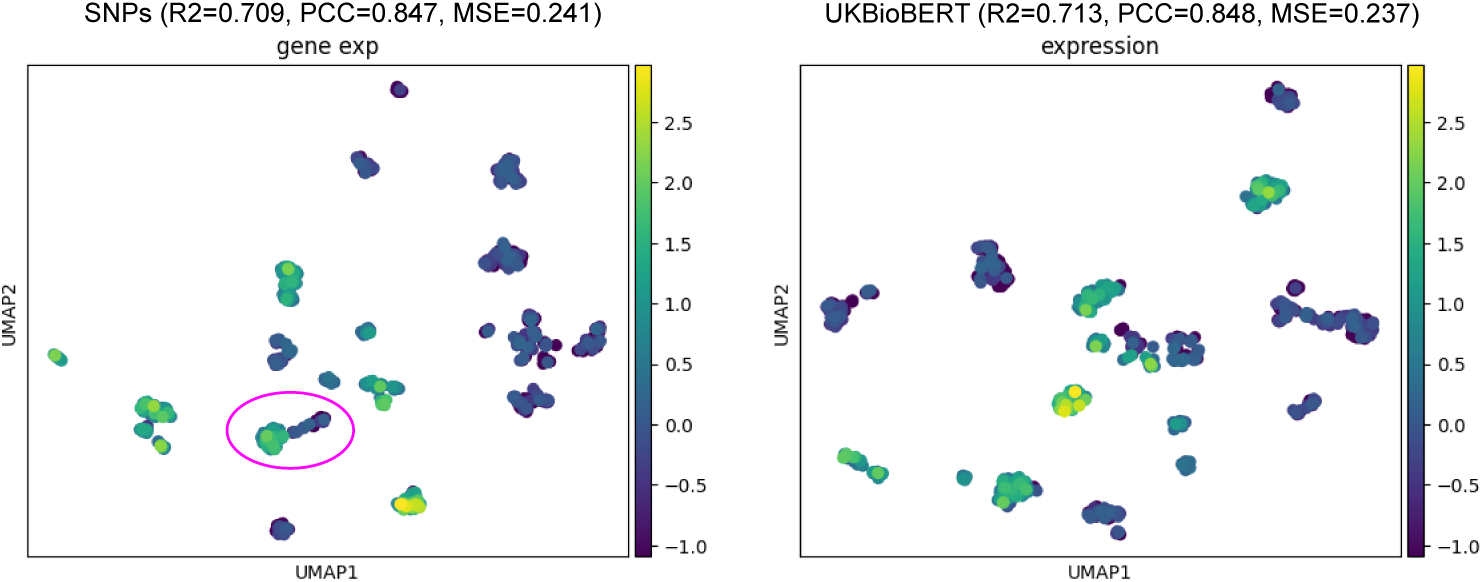
UMAP visualization of individuals based on SNPs (left panel) or gene embeddings from UKBioBERT (right panel) colored by expression levels of ZFP57. The performances of gene expression prediction based on different input data are annotated with figure legends. The confusing area clustered by UMAP with SNPs is highlighted with a pink circle.

**Extended Data Fig. 8.**
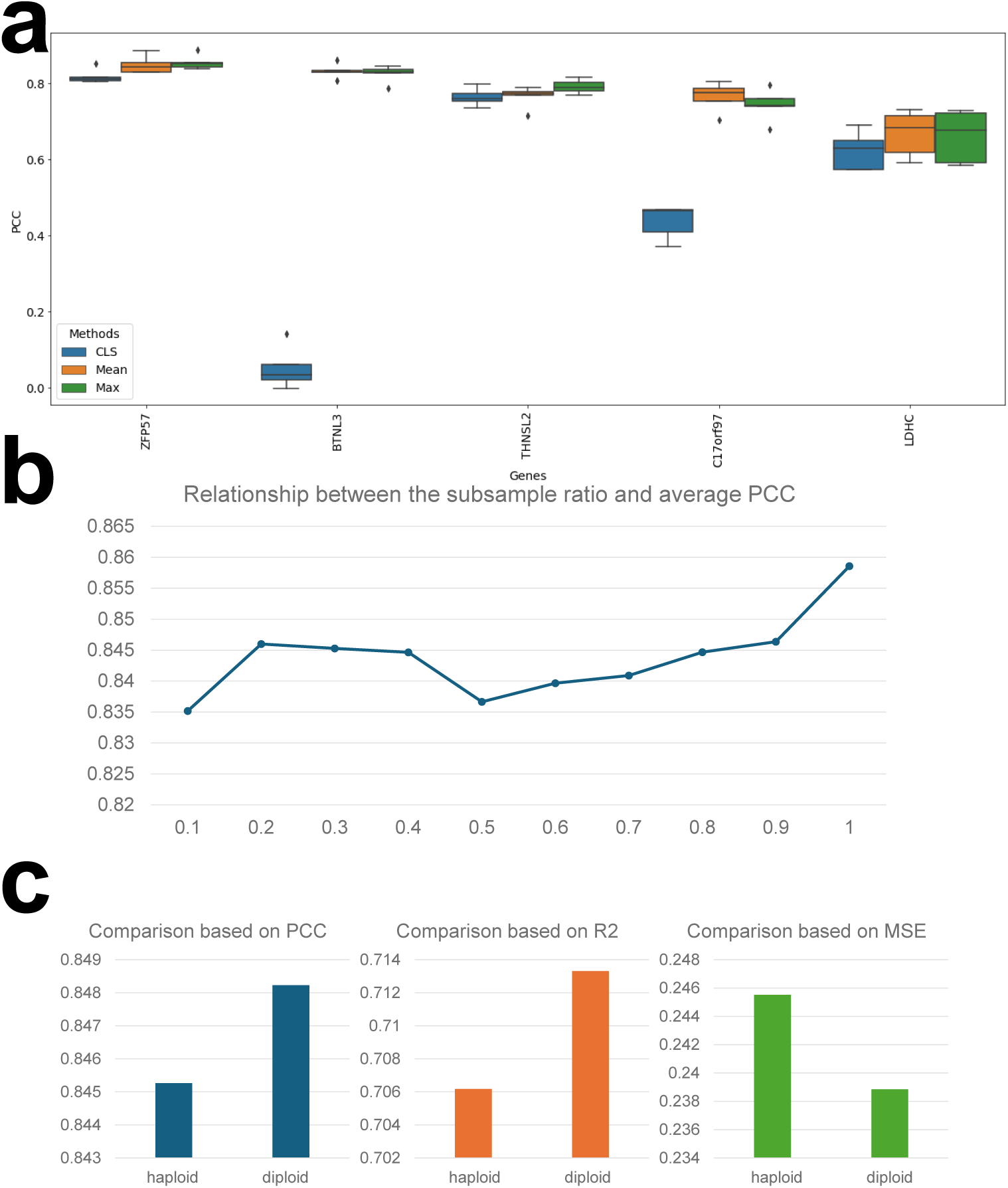
Sensitivity analysis of UKBioBERT. (a) Comparisons of different pooling settings for gene expression prediction. (b) Relationship between the subsample ratio and average PCC to test data-efficient learning. (c) Comparison between using haploid information for gene expression prediction and diploid information for gene expression prediction. The scores of metrics are computed based on averaging five folds.

**Extended Data Fig. 9.**
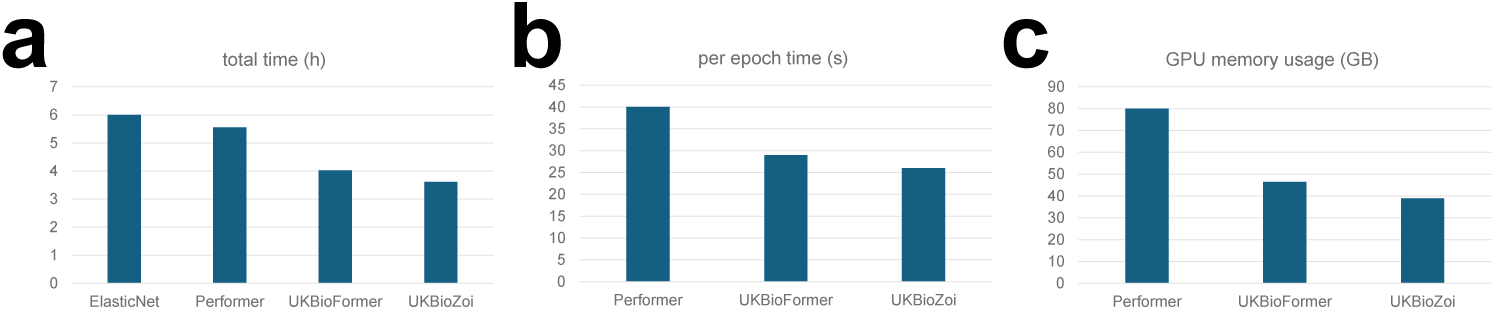
Time and memory usage of fine-tuning process. (a) Comparison of total training-testing time for selected models based on the five-fold cross validation with a single gene. (b) Comparison of batch-level training-testing time for selected models based on the five-fold cross validation with a single gene. (c) Comparison of GPU consumption for selected models based on the five-fold cross validation with a single gene.

**Extended Data Fig. 10.**
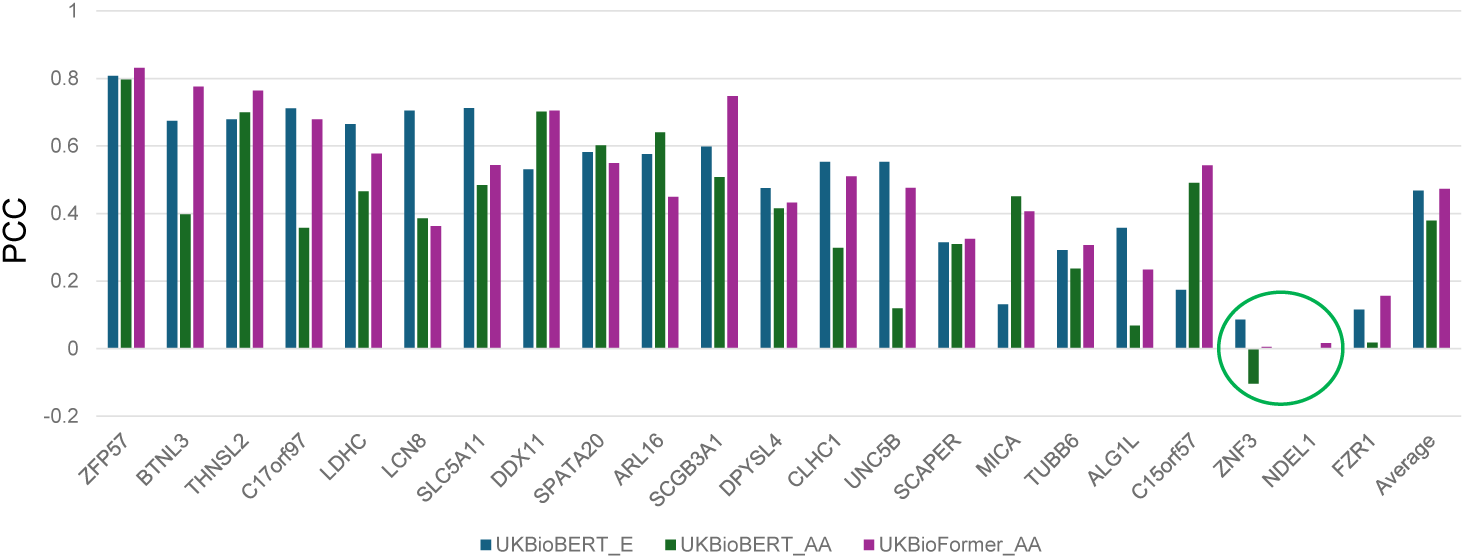
Comparison of selected models (UKBioBERT and UKBioFormer) for cross-cohort gene expression prediction. The sample with negative correlation predicted by UKBioBERT for the AA group is highlighted by a green circle, while UKBioFormer did not face such problem.

**Extended Data Fig. 11.**
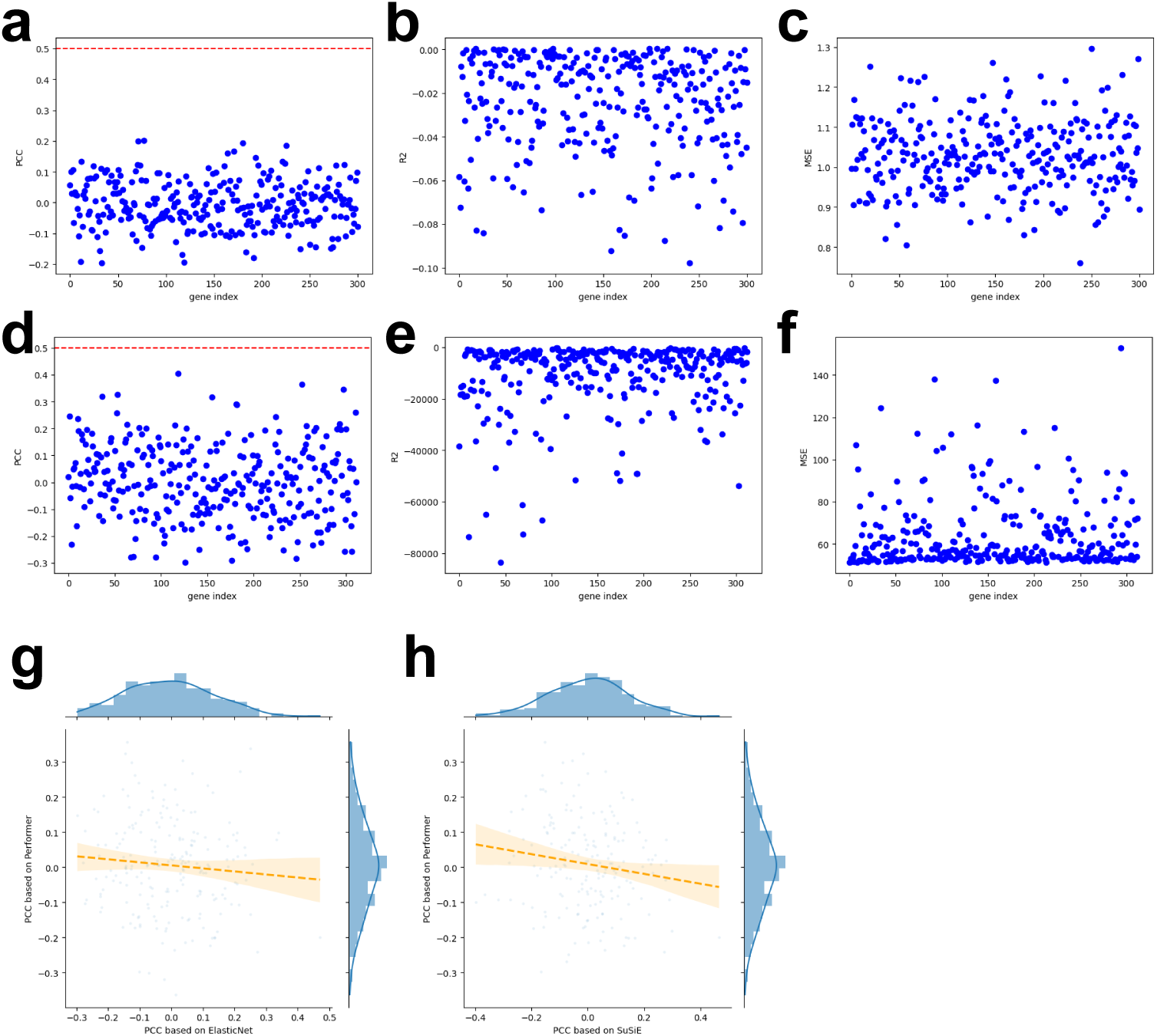
Performance of UKBioFormer with joint training 300 genes from GTEx data and ROSMAP data. (a)-(c) represents the joint training result of GTEx data based on whole blood. (d)-(f) represents the result based on GTEx brain cortex data training and ROSMAP brain cortex data testing. (a) PCC score of each gene. (b) R2 score of each gene. (c) TSE score of each gene. (d) PCC score of each gene. (e) R2 score of each gene. (f) MSE score of each gene. (g) Relationship between the PCC computed based on the results from ElasticNet and the PCC computed based on the results from Performer. (h) Relationship between the PCC computed based on the results from SuSiE and the PCC computed based on the results from Performer.

**Extended Data Fig. 12.**
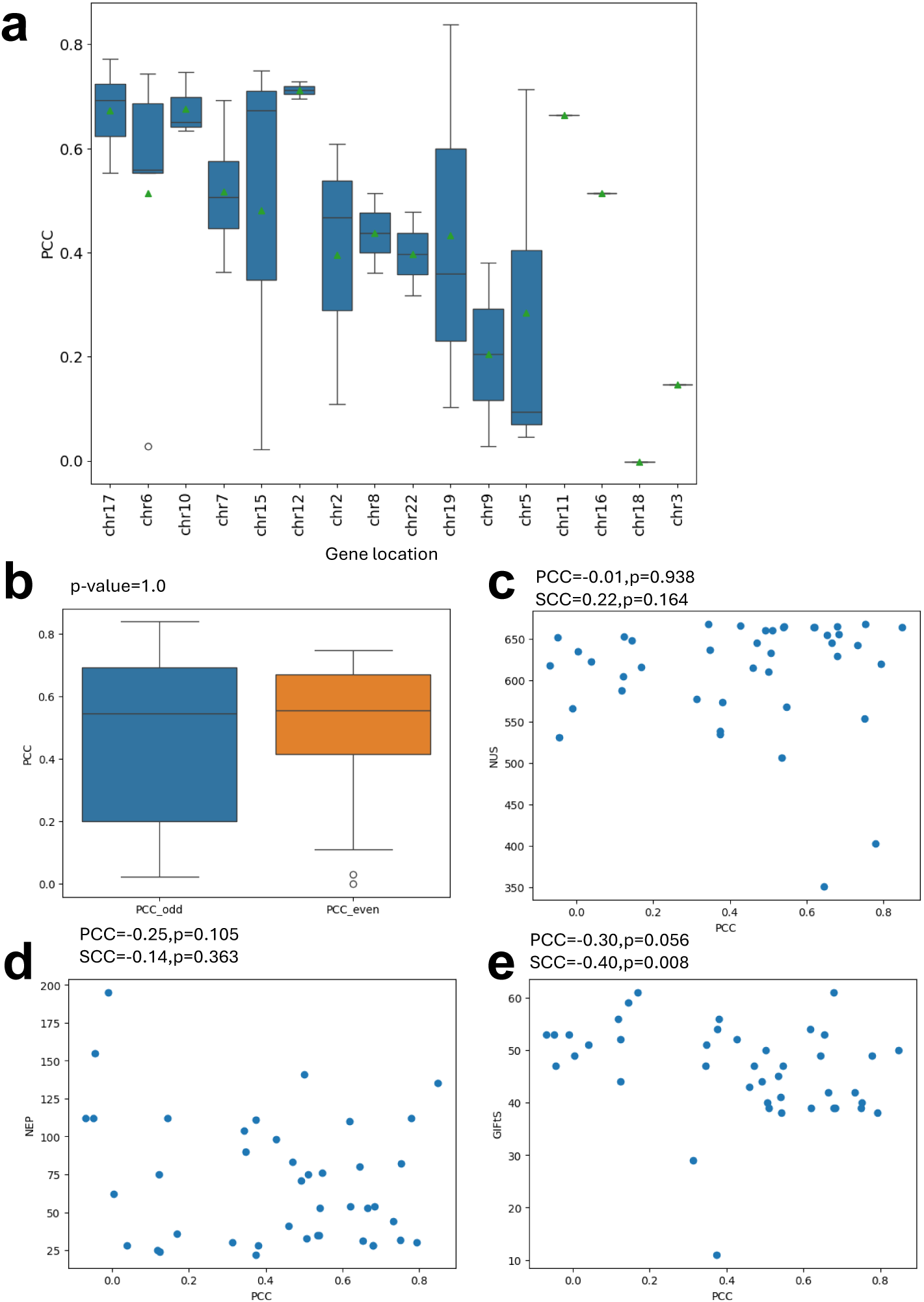
Analyzing factors affecting gene predictability. Panels (b)-(e) are annotated with related statistics. (a) Boxplot used to represent the distribution difference of PCCs from genes with different chromosome location. For chromosomes chr11, chr16, chr18, and chr3, they only contains one gene for testing. (b) Boxplot used to represent the distribution difference of PCCs from genes with different chromosome location grouped by odd chromosome and even chromosome. (c) Relationship between NUS and PCCs. (d) Relationship between NEP and PCCs. (e) Relationship between GIFts and PCCs.

**Extended Data Fig. 13.**
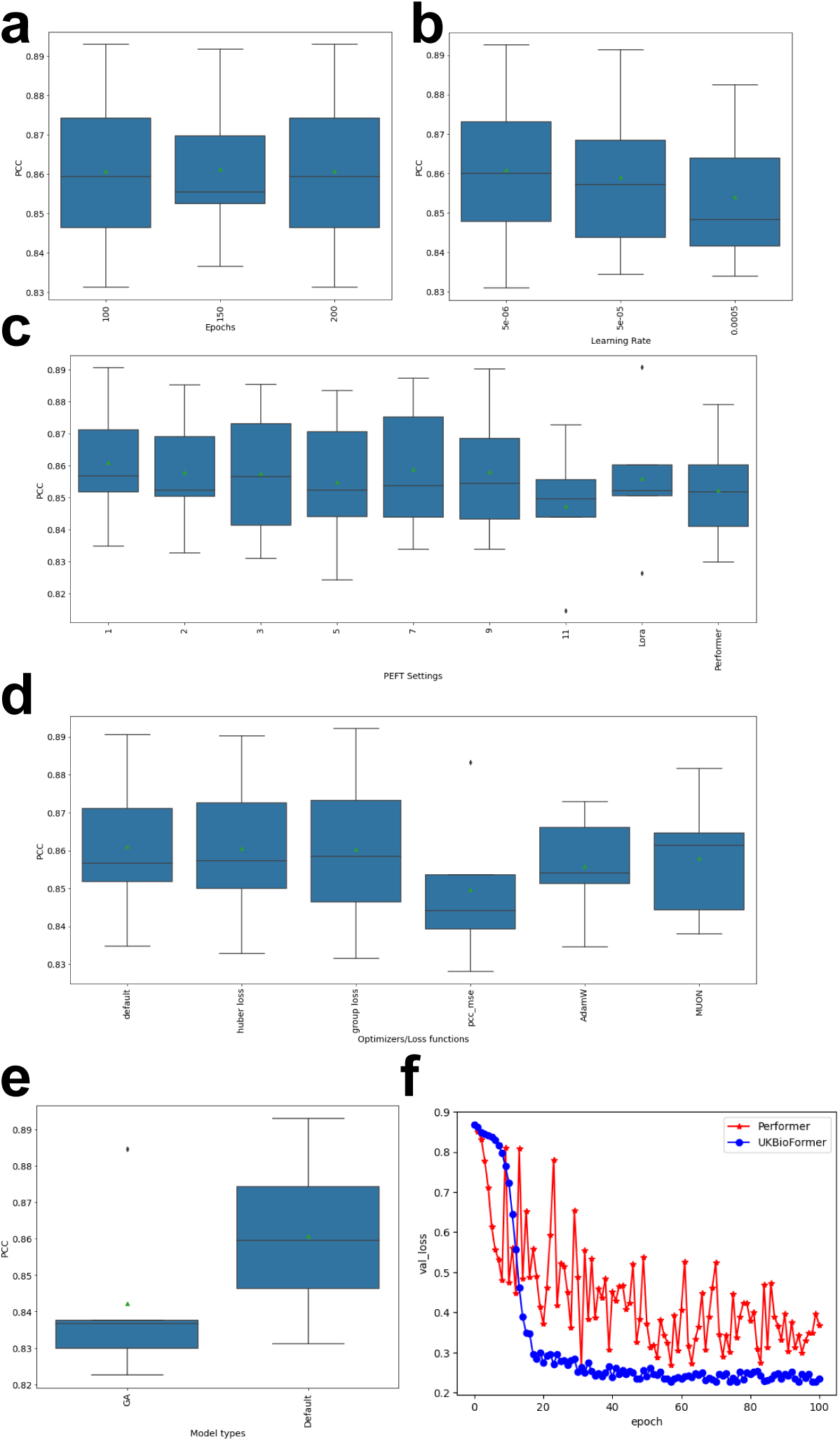
Results of ablation tests and hyper-parameter tuning based on XFormer models. We present the five-fold cross-validation result for each setting. (a) Relationship between PCCs and epochs. (b) Relationship between PCCs and learning rates. (c) Relationship between PCCs and different fine-tuning settings. (d) Relationship between PCCs and optimizers/loss functions. (e) Relationship between PCCs and gradient accumulation (GA) usage. (f) Relationship between training epochs and corresponding validation loss (val loss) for Performer and UKBioFormer.

**Extended Data Fig. 14.**
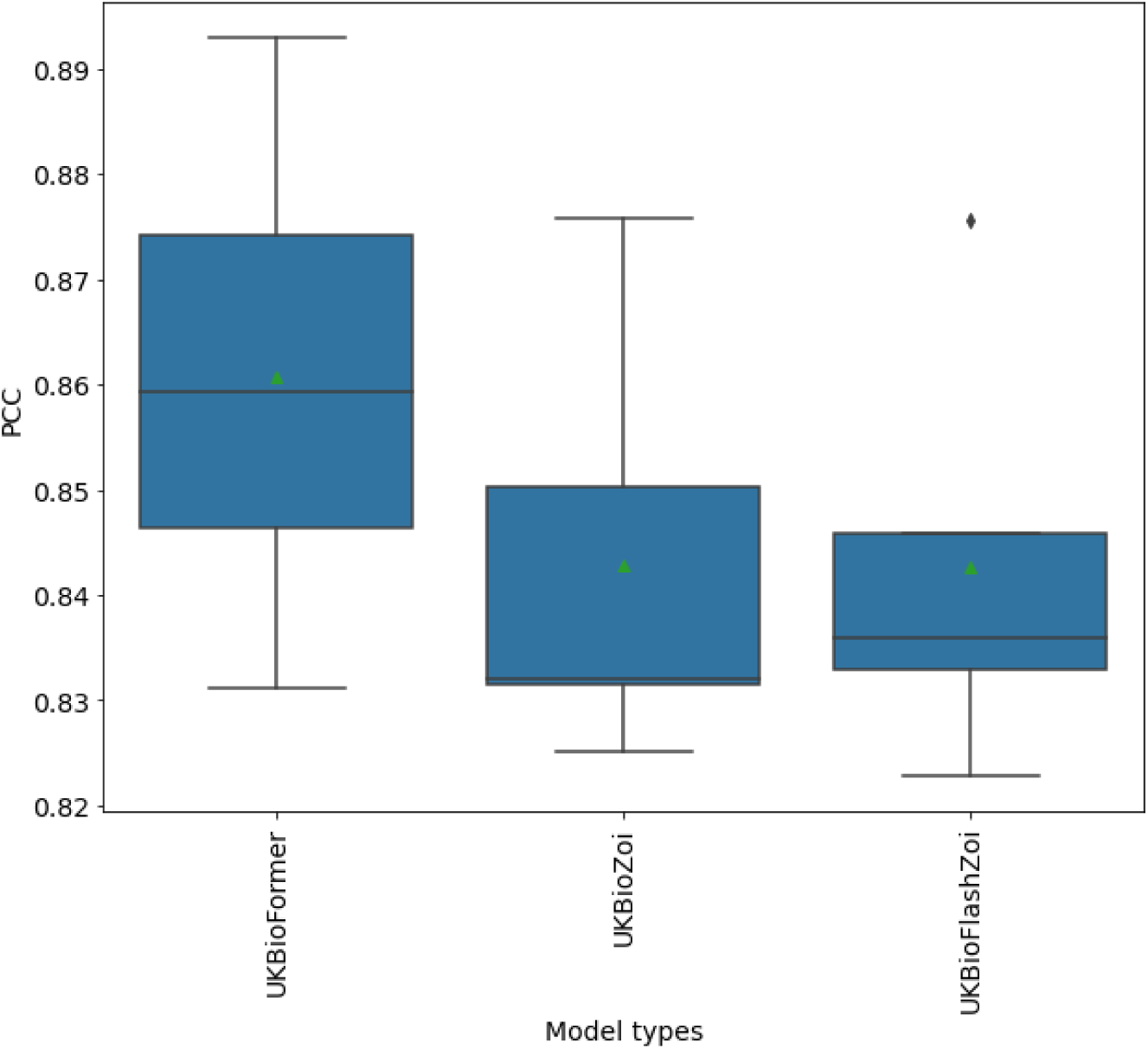
Comparison among UKBioFormer, UKBioZoi, and UKBioFlashZoi based on PCC.

**Extended Data Fig. 15.**
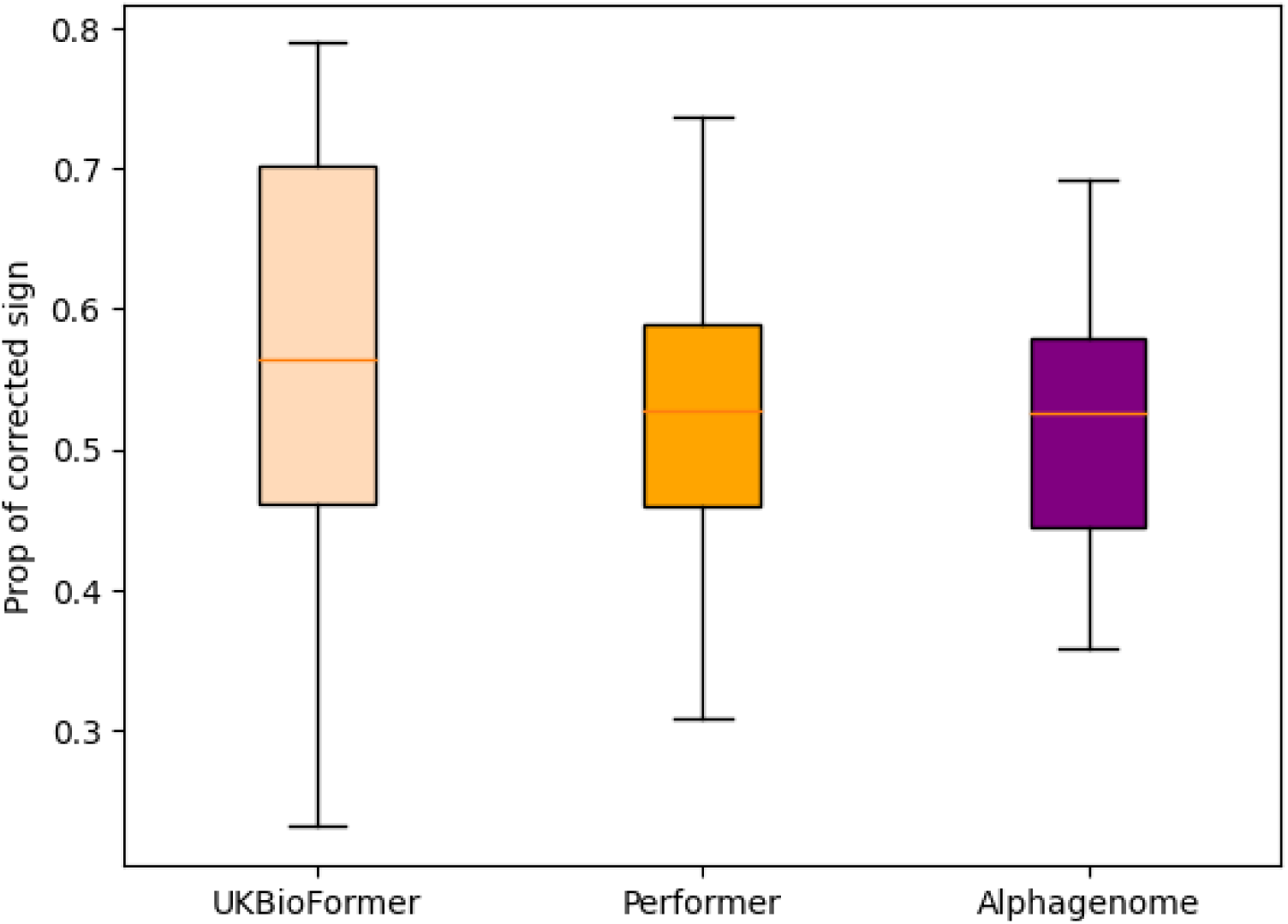
The comparisons among UKBioFormer prediction results, Performer prediction results, and Alphagenome prediction results for eQTL identification.

**Extended Data Fig. 16.**
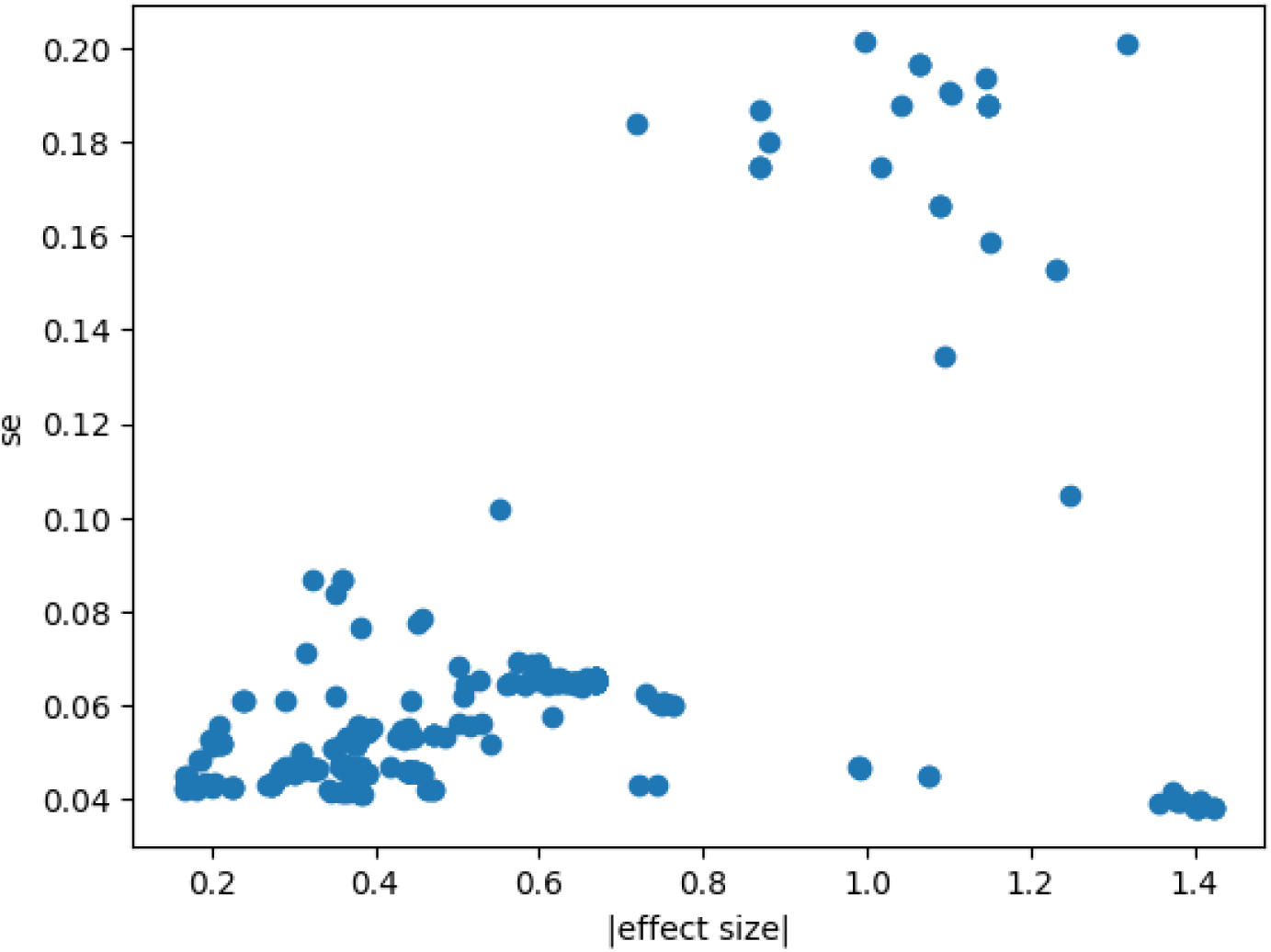
Relationship between absolute effect size and standard error.

**Extended Data Fig. 17.**
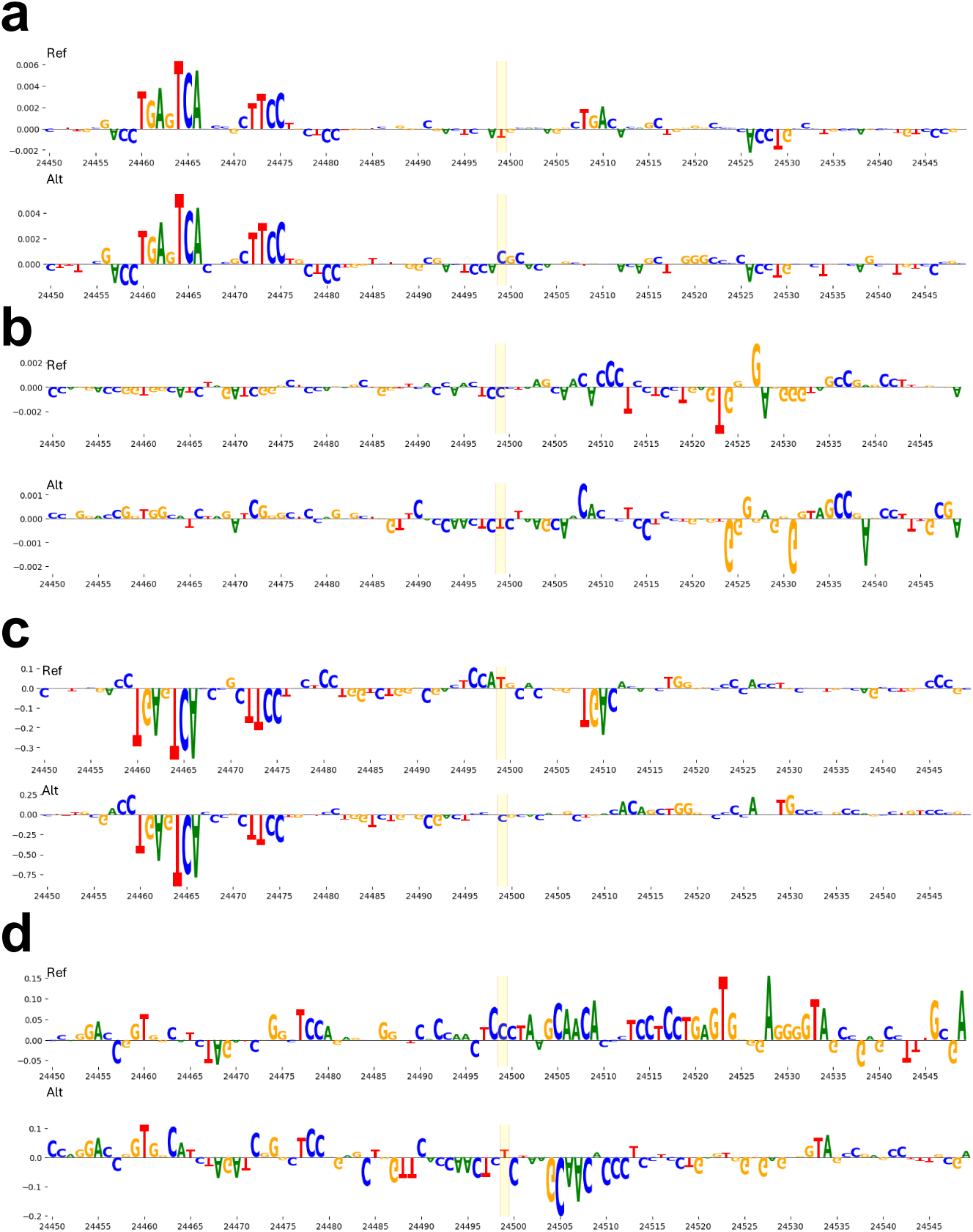
Variant importance estimated based on Enformer (shown in (a) and (b)) and Performer (shown in (c) and (d)). Figures (a) and (c) represent the results of variant rs9910080. Figures (b) and (d) represent the results of variant rs9903086.

